# Spatially resolved niche and tumor microenvironmental alterations in gastric cancer peritoneal metastases

**DOI:** 10.1101/2024.03.21.585686

**Authors:** Joseph J. Zhao, Johnny Chin-Ann Ong, Supriya Srivatsava, Daryl Kai Ann Chia, Haoran Ma, Kiekyon Huang, Taotao Sheng, Kalpana Ramnarayanan, Xuewen Ong, Su Ting Tay, Takeshi Hagihara, Angie Lay Keng Tan, Melissa Ching Ching Teo, Qiu Xuan Tan, Gillian Ng, Joey Wee-Shan Tan, Matthew Chau Hsien Ng, Yong Xiang Gwee, Robert Walsh, Asim Shabbir, Guowei Kim, Yvonne Tay, Zhisheng Her, Giuseppe Leoncini, Bin Tean Teh, Jing Han Hong, Ryan Yong Kiat Tay, Chong Boon Teo, Mark P. G. Dings, Maarten Bijlsma, Jeffrey Huey Yew Lum, Jia Hao Law, Filippo Pietrantonio, Steven M. Blum, Hanneke van Laarhoven, Samuel J. Klempner, Wei Peng Yong, Jimmy Bok Yan So, Qingfeng Chen, Patrick Tan, Raghav Sundar

## Abstract

Peritoneal metastases (PM) in gastric cancer (GC) portend a poor prognosis, yet our understanding of tumor microenvironmental (TME) characteristics associated with GCPM remain limited. Here, we analyzed intrinsic genomic alterations and transcriptomic programs predictive of GCPM in a prospective cohort of 248 patients, identifying *CDH1*, *PIGR*, and *ELF3* mutations as predictors. By inspecting the spatial dynamics of the TME, we find that tumor compartment infiltration of pro-tumorigenic cell types such as inflammatory cancer-associated fibroblasts (CAFs) predict peritoneal recurrence. Next, in a cross-sectional study of 205 samples from 55 patients, distinct pathways and immune compositions in GCPM relative to liver metastases highlight the TME’s significance in transcoelomic metastases. Notably, several putative therapeutic targets exhibited distinct expression patterns between PTs and PMs. We also observed increased immune infiltration in GCPMs treated with systemic immunotherapy and intraperitoneal chemotherapy. Our findings highlight transcriptomic variations and niche reprogramming in the GCPM peritoneal environment, revealing roles of myeloid dendritic cells, effector memory CD8+ T cells, and CAFs in metastatic progression.

**Statement of significance:** Comprehensive molecular profiling of gastric cancer primary and peritoneal tumors unveils crucial insights into the distinct molecular and immune landscape of peritoneal metastases. Identifying predictive markers and therapeutic targets emphasizes the significance of tumor microenvironment alterations in guiding future therapies for gastric cancer peritoneal metastasis.

## Introduction

The peritoneum is a common site of metastases in gastric cancer (GC). Peritoneal metastasis (PM) is associated with poor prognosis^1^ and significant morbidity. PM may cause intestinal obstruction, malnutrition and impaired bowel function, as well as malignant ascites.^2^ In contrast to liver or lung metastases where tumor cells are required to enter the blood circulation (systemic dissemination), direct seeding into the peritoneal cavity (transcoelomic metastasis) is presumed to be the most common route of PM.^3^ Although granular mechanistic insights are lacking, successful peritoneal dissemination appears to require a series of biological traits: anchorage-independent survival, adherence and implantation to serosal surfaces, creation of a suitable microenvironment, neo-angiogenesis, and immune evasion.^3,4^

The proclivity of GCs to metastasize to the peritoneum has been hypothesized to occur due the latter’s immunosuppressive microenvironment, analogous to other immune niches such as the brain and bone.^5,6^ The conditioning of a spatially distant location prior to colonization – the pre-metastatic niche – has been speculated to occur through education from the primary tumor (PT).^6,7^ For transcoelomic metastases to occur, several components of the preparation of a pre-metastatic niche have been described and hypothesized. These include secretion of PT-derived factors that contribute to, immune suppression, organ-specificity or organotropism, lymph-angiogenesis, and reprogramming of metabolic and epigenetic pathways.^6,8^ However, these hypotheses have not been firmly established in PM, largely because of the lack of studies performed directly on peritoneal tissue and peritoneal-resident samples and limited immunocompetent model systems. While studies on malignant ascites in GC^9–11^ have provided an initial characterization of PM, analyses restricted to cells and substrates secreted into the free-floating ascites preclude holistic comprehension of PM biology such as the *in situ* PM tumor microenvironment (TME). The development of malignant ascites is also likely a late biological event in the evolution of PM, and studies restricted to this sample source may limit insight into early dissemination mechanisms.^12^ Similarities and differences between gastric PTs and the peritoneal niche thus remain poorly understood.

Recently, laparoscopic approaches to evaluate and deliver peritoneal-directed therapy has allowed for paired and repeated sampling of PM. As such, these elusive tumors which were historically difficult to access and study in detail are now being routinely collected for translational analyses. In this study, we sought to understand genomic, transcriptomic and TME features that determine and facilitate peritoneal organotropism from primary GCs. By performing one of the largest multi-omic molecular characterization of GCPM, we unveiled biological principles underpinning transcoelomic metastasis in GC.

## Results

### Cohort summary and overview

To identify intrinsic genomic alterations and transcriptomic programs associated with PM, we analyzed 276 samples from a prospective cohort of primary GCs from 248 patients with longitudinal PM outcomes after primary resection. Thereafter, based on hypotheses generated from the prospective cohort, we analyzed a separate cross-sectional cohort of metastatic GCs comprising 205 samples from 55 patients with PM. Peritoneal tissue samples from GC patients were predominantly collected during diagnostic staging laparoscopy and/or during peritoneally-directed clinical trials such as intraperitoneal catheter-based chemotherapy and pressurized intraperitoneal aerosol chemotherapy (PIPAC).^13^

The samples were profiled with three assays: whole exome sequencing (WES), whole transcriptome sequencing (WTS) and digital spatial profiling of independent tumor and stroma compartments (DSP, GeoMx platform, Nanostring Technologies, Inc). For tissue that was flash-frozen, bulk WES and WTS was performed. For formalin-fixed paraffin embedded (FFPE) tissue, spatial profiling was performed using the NanoString GeoMx platform [**Fig. 1a** and **Fig. 1b**]. Synchronous PM was defined as PM diagnosed within 6 months of cancer diagnosis while metachronous PM was defined as PM diagnosed after 6 months of cancer diagnosis. Of the 303 patients in total, 51 (16.8 %) patients had metachronous PM and 58 (19.1%) patients had synchronous PM. Median age was 69.0 (IQR 60.0 – 76.0) years and majority of patients were male (57.2%), Chinese (85.9%). Lauren’s histology classification of intestinal subtype was reported in 47.1% and diffuse subtype in 27.5% of PT [**Supplementary Table 1a**]. Median follow-up was 81.9 months. Samples analyzed included PTs from the stomach (n=268), adjacent normal tissue to the PT (n=44), PM (n=108) and adjacent normal peritoneal tissue (n=61). An overview of paired samples is presented in **Supplementary Table 1b**.

**Fig. 1.**
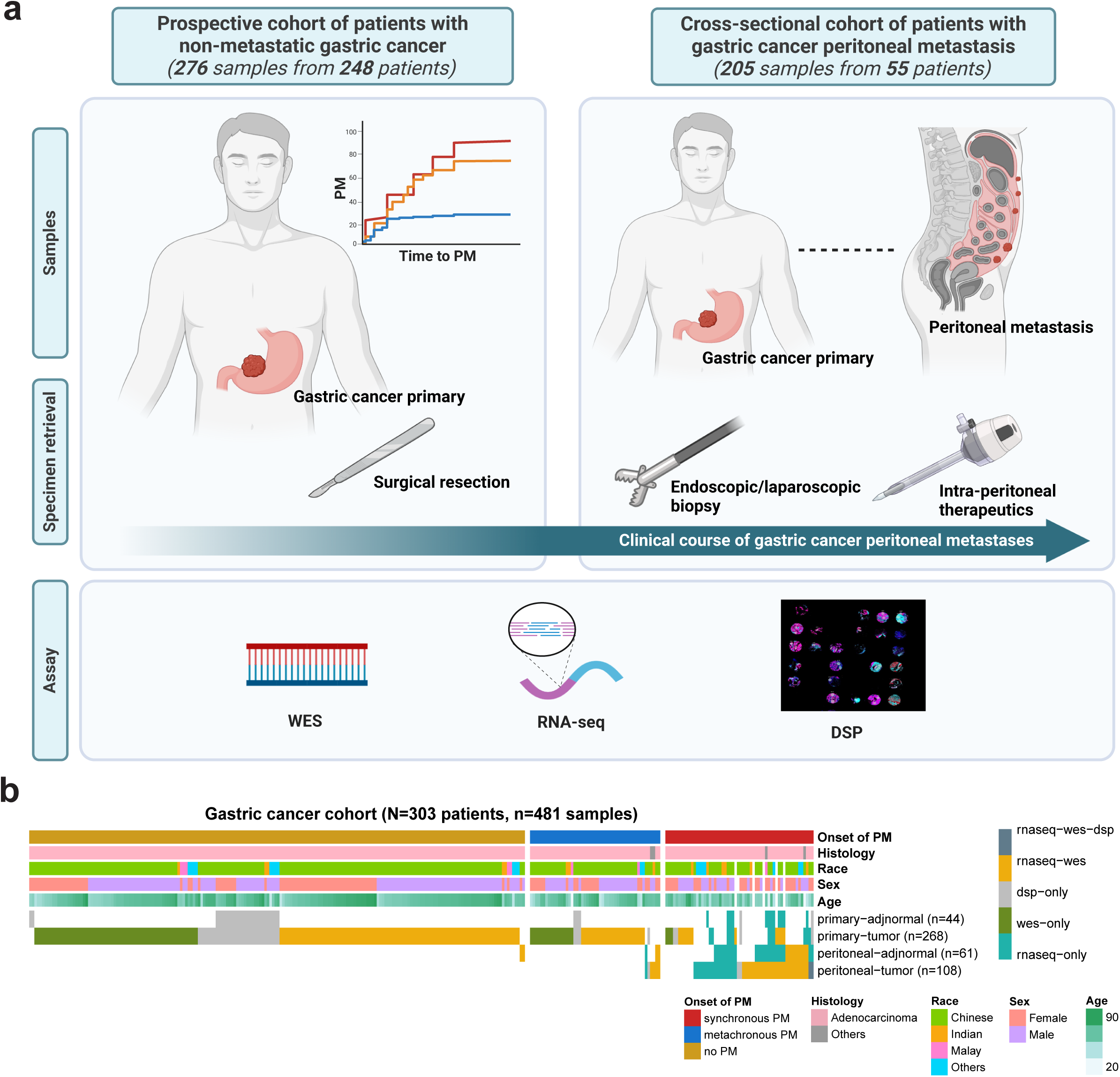
Study and cohort overview. **a**. Schematic illustration of study overview. This illustration was created using BioRender.com. **b.** Cohort overview of participants with gastric cancer, describing the distribution of retrieved samples and profiling assay. Each patient may have more than one sample per sequencing modality. The above sample count excludes 11 benign normal peritoneal tissue from patients with benign conditions and 20 samples of primary tumor with liver metastasis/liver metastases utilized in downstream analyses. Abbreviations: N=, number of patients; n=, number of samples; WES, whole exome sequencing; RNA-seq, RNA sequencing; DSP, digital spatial profiling; PM, peritoneal metastasis; adjnormal, adjacent normal.

### Molecular aberrations and pathways in primary GC tumors predictive of peritoneal metastasis

In the prospective cohort, fifty-six patients (22.6%) developed PM after a median follow-up duration of 87.0 months and had poorer OS compared to patients without metastasis (log-rank, p<0.001) [**Supplementary Fig. 2a**]. Adjusted for competing events of death, patients with bulkier and more advanced tumors were at increased hazard of PM (T-stage and N-stage stratified analysis, log-rank, p=3.81e-07 and p=4.94e-06 respectively). Consistent with known clinical outcomes, TCGA GS subtype^14^ (log-rank, p=0.00115) and mesenchymal subtype GC^15^ were associated with a higher tendency to develop PM (log-rank, p=0.00214). Participants with Lauren’s intestinal subtype classification were at a lower hazard of PM compared to the diffuse/mixed/NOS subtype (log-rank, p=0.00885). WTS derived fibrotic or immune-enriched/fibrotic TME subtypes (F or IE/F) were also at a higher risk of peritoneal recurrence (p=0.00267) [**Fig. 2a**].

**Fig. 2.**
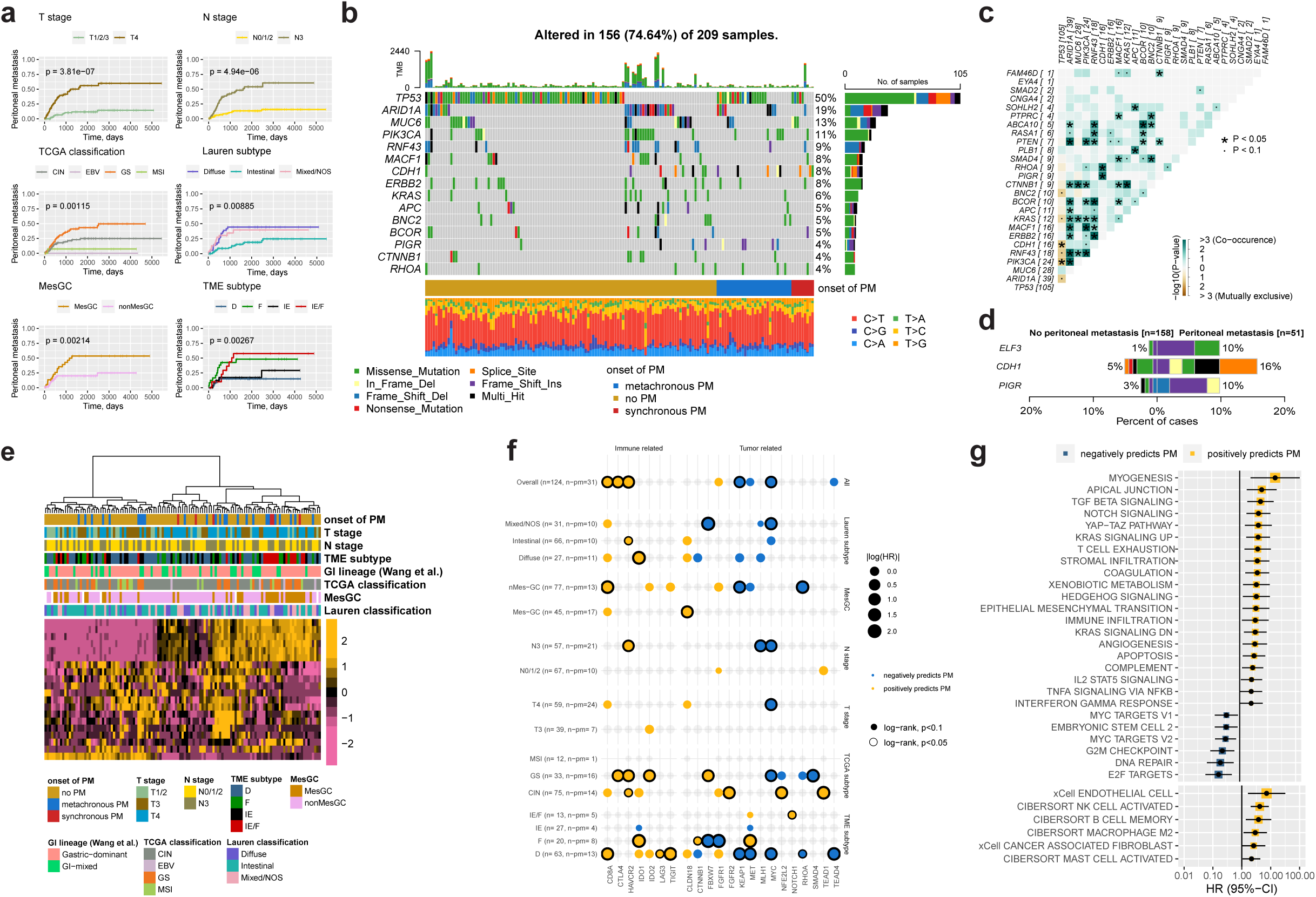
Oncogenes and pathways driving peritoneal metastasis in gastric cancer. **a**. KM curves of PM stratified by T & N-stage, Lauren, TCGA classification, MesGC and TME subtype. P-values were retrieved from the Fine-Gray subdistribution hazard model. **b**. Oncoplot showing driver mutations identified in WES of GC PT samples in the prospective cohort. Genes shown here are driver mutations previously reported by Bass *et al*^14^ **c**. Somatic interactions between co-occurring and mutually exclusive driver mutations. P-values were retrieved from pairwise two-sided Fisher’s Exact test. **d**. Mutations significantly associated with GCPM. Colors representing the type of mutation follow the legend in Fig. 2a. **e.** Heatmap of gene expression profiles between GC PT samples with and without PM (genes were selected if standard deviation of log2(FPKM) values were greater than 5). **f.** Dot plot of selected genes associated with PM in the overall cohort and respective subgroups. P-values were derived from a log-rank test. **g**. Forest plot of pathways and immune cell types predictive of PM. Abbreviations: KM, Kaplan-Meier; TCGA, The Cancer Genome Atlas; GC, gastric cancer; Mes, mesenchymal subtype; nMes, non-mesenchymal subtype; TME, tumor microenvironment; F, fibrotic; IE, immune-enriched; D, depleted; NOS, none otherwise specified; CIN, chromosomal instability; GS genomically stable; MSI, microsatellite instability; EBV, Epstein-Barr virus; WES, whole exome sequencing; PT, primary tumor; HR, hazard ratio; CI, confidence interval; n=, number of patients; n-pm, number of patients with peritoneal metastasis; PM, peritoneal metastasis.

We performed WES on 209 PT GC samples (average coverage 129X for tumor samples, 70X coverage for matched normal/blood controls). Recurrent oncogenic alterations in the overall cohort are shown in **Fig. 2b**. Among patients who developed PM, *TP53* was the most commonly altered driver mutation (25/51, 49.0%), followed by *ARID1A* (9/51, 17.6%), *CDH1* (8/51, 15.7%), *PIK3CA* (6/51, 11.8%) [**Supplementary Fig. 2b, c**]. Evaluation of somatic interactions found that *CDH1* and *TP53* mutations were mutually exclusive. Conversely, *CDH1* mutations were found to significantly co-occur with *PIGR* and *RHOA* mutations. *ARID1A* mutations were also found to significantly co-occur with multiple other mutations such as *PTEN*, *PIK3CA*, *ERBB2*, and *KRAS* [**Fig. 2c**].

Comparing the PTs of patients with PM against those PTs without PM in the prospective cohort, *ELF3*, *CDH1*, *PIGR* were significantly associated with PM [**Fig. 2d**]. Subgroup analyses further identified additional driver genes associated with PM in GS patients. Amongst GS tumors, *ARID1A* mutations (OR=6.34, P-value=0.018) were significantly associated with PM compared to patients with GS tumors with no metastasis [**Supplementary Fig. 2d**]. These aberrations were largely consistent with previously reported driver genes.^11,16^ Bulk RNA-seq (WTS) was performed on 124 PT samples (59.3%). Unsupervised, hierarchical clustering demonstrated distinct gene expression profiles between patients with and without PM [**Fig. 2e**]. Patients with high expression of immune-related genes such as *TIM3* (*HAVCR2*) and *CD8A* were associated with an increased hazard of PM across multiple subgroups and in the overall cohort; conversely, high *CLDN18* expression was associated with PM only in high-risk subgroups such as MesGC (HR=4.384, log-rank p=0.0087) [**Fig. 2f**]. Other comparisons are detailed in **Fig. 2f**.

Single sample Gene Set Enrichment Analysis (ssGSEA) pathway analysis showed that PT samples with a greater hazard of developing PM exhibited high (upper third vs lower third) stromal infiltration, epithelial mesenchymal transition (EMT), inflammatory pathways (transforming growth factor-β [TGF-β] signalling, IL2-STAT5, interferon gamma response [IFNCγ], tumor necrosis factor-α [TNF-α] signalling) and T cell exhaustion enrichment scores. Embryonic stem cell signature scores were inversely associated with PM in GC (HR=0.33, log-rank p=0.015), discordant with previous hypotheses on the role of embryonic stem cells in GC transcoelomic metastasis.^17^ Samples with high MYC enrichment scores were found to have a lower hazard for PM [**Fig. 2g**]. Comparison of CIBERSORT and xCell enumerated immune cell expressions found that increased cancer associated fibroblasts (CAFs) (HR=2.75, log-rank P-value=0.040) and M2 macrophages (HR=2.91, log-rank P-value=0.019) in PT were associated with a greater hazard of PM [**Fig. 2g**].

Spatial transcriptomics was conducted on 67 samples (37 PT and 30 primary adjacent normal) from 39 patients [**Fig. 3a**]. After a median follow up of 46.0 months, 5 (16.7%) patients were found to have peritoneal recurrence. Grossly, we found that PT-tumor compartments from patients with peritoneal recurrence were transcriptomically closer to PT-stromal compartments after dimensionol reduction of gene expression profiles [**Fig. 3b**]. EMT, stromal and immune infiltration pathways were enriched in patients with peritoneal recurrence [**Fig. 3c**]. In 27 patients (n=23 patients without peritoneal recurrence, n=4 patients with peritoneal recurrence) where paired stromal and tumor compartments within PT samples were available, we found a trend towards higher enrichment of fibroblasts, CD8 T cells and mast cells within tumor compartments in patients with peritoneal recurrence. Conversely, NK cells, plasma cells, Tregs, mast cells and macrophages were relatively enriched in stromal regions of PT in patients with peritoneal recurrence [**Fig 3d**]. Although macrophage expression was found to be lower within tumor compartments in patients with peritoneal recurrence, the TAM M2/M1 ratio signature suggested by Jiang *et al.*^18^ was found to be higher in PT-stromal compartments and enriched in PT-tumor compartments amongst participants with peritoneal recurrence [**Fig. 3e**] Further, we found that the IL iCAF signature by Kieffer *et al.*^19^ was significantly higher in tumor compartments of patients with PM. Taken collectively, our findings allude to the possibility that tumor compartment infiltration of these tumorigenic cell types may be associated with peritoneal recurrence. Collectively, analysis of PT samples from our prospective cohort suggest PT TME-specific traits are associated with GCPM.

**Fig. 3.**
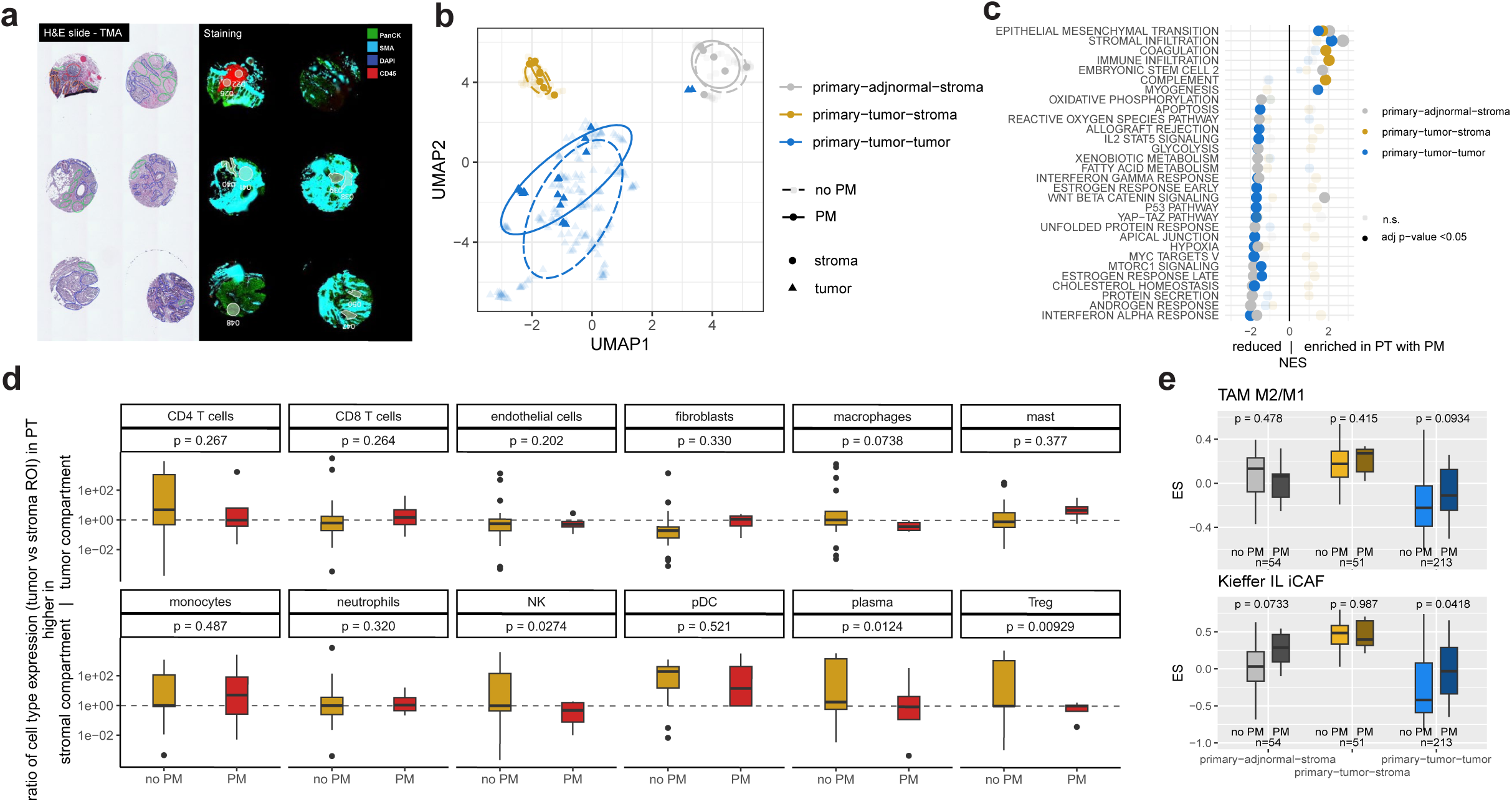
Spatially resolved microenvironmental changes associated with peritoneal recurrence. **a.** H&E slide, staining of TMA for DSP. **b.** UMAP of sequenced ROIs, stratified by peritoneal recurrence. **c.** GSEA comparisons between patients with and without peritoneal recurrence of pathways within each ROI. **d.** Ratio of *SpatialDecon* numerated cell type expression between PT tumor and stroma. P-values were retrieved with an unpaired T-test. **e.** Differences in GSVA enrichment scores of IL iCAFs (Kieffer *et al.*) and M2/M1 TAM (Jiang *et al*.) against PM outcomes across ROIs. Abbreviations: ROI, regions of interest; DSP, digital spatial profiling; TMA, tumor microarray; GSEA, gene set enrichment analysis; PM; NES, normalized enrichment score; PM, peritoneal metastasis; TAM, tumor associated macrophages; iCAF, inflammatory cancer associated fibroblasts; ES, enrichment score; adjnormal, adjacent normal; adj, adjusted.

### Evolution of transcriptional programs and tumor microenvironment in GCPM

Next, we explored PT-PM differences to study pathways associated with transcoelomic metastasis. A cross-sectional cohort of GCPM samples was integrated with the upstream prospective cohort [**Supplementary Fig. 4a, b**]. To compare differences in metastases to the peritoneum compared metastases to other sites, we harnessed a cohort of patients with gastroesophageal junction cancers and liver metastases^20^.

Unsupervised UMAP clustering of GC WTS samples demonstrated distinctions between three groups - PT without PM, PT with synchronous or metachronous PM, and PM samples. Pseudotime analyses of these samples demonstrated a trajectory from PT without PM, progressing to PT that develop PM and eventually the PMs themselves. Transcriptomic trajectories towards liver metastases were found to be divergent and distinct from PM [**Fig. 4a**]. Across PT without PM (n=94), PT with PM (n=49) and PM (n=100) WTS samples, pathways previously identified in the prospective cohort to be associated with PM development in GC PT samples such as EMT, angiogenesis and the *YAP-TAZ* pathway were shown to be further overexpressed in PM samples, reinforcing its likely role in PM development. Immune composition of CAFs (iCAFs/myCAFs), M2 macrophages, monocytes, activated NK cells and dendritic cells were notably increased in PMs compared to PTs. Conversely, naïve CD4+ T cells were reduced in PM [**Fig. 4b**].

**Fig. 4.**
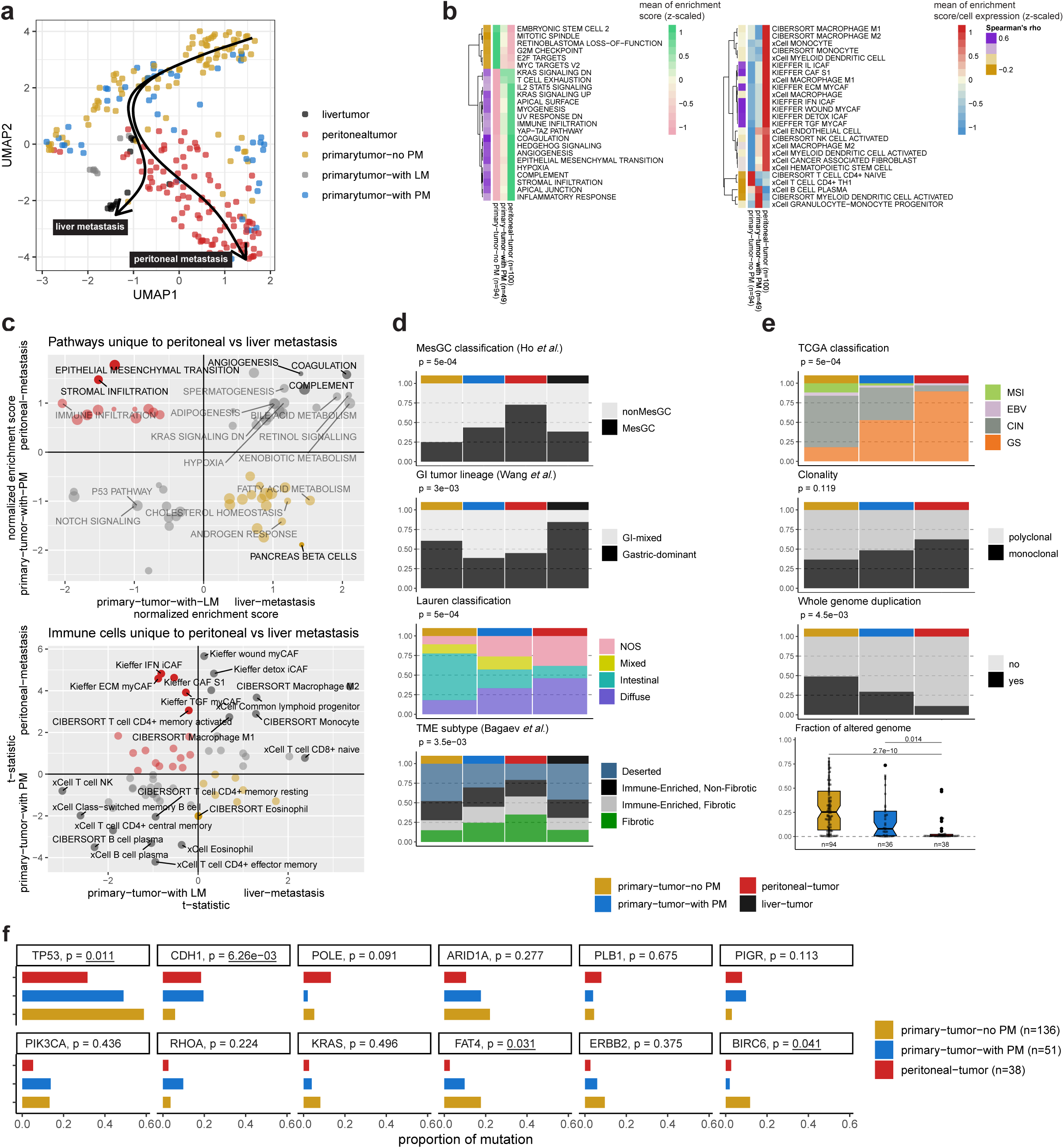
Evolution of microenvironmental and oncogenic drivers in gastric cancer peritoneal metastasis. **a.** UMAP of bulk RNA-sequenced GC tumor samples. Arrows represent trajectories derived from pseudotime analyses. **b.** Heatmap of ssGSEA mean enrichment scores per sample type for selected pathways involved in PM and mean CIBERSORT/xCell enumerated cell expressions in bulk RNA-seq samples. **c**. Scatter plot of NES from pairwise GSEA and t-statistics of CIBERSORT/xCell immune cell types between PT and PM/LM. **d.** Proportions of mesenchymal-type gastric cancer, GI tumor lineages, TME subtypes and proportion of primary tumor Lauren’s classification across PT without PM, PT with PM and PM. **e.** Proportions of TCGA classification, clonality, WGD and fraction of altered genome across PT without PM, PT with PM and PM. P-values for **Fig. 4d & 4e** were retrieved using the Fisher’s exact test. Raw values for **Fig. 4d & 4e** may be found in **Supplementary Table 4a**. **f.** Comparisons of underlying driver mutations across PT without PM, PT with PM and PM. P-values were retrieved with the Fisher’s exact test. Abbreviations: GC, gastric cancer; WGD, whole-genome duplication; PM, peritoneal metastasis; PT, primary tumor; LM, liver metastasis; TME, tumor microenvironment; GSEA; Gene Set Enrichment Analysis; GSVA, gene set variation analysis.

Pairwise pathway analyses between GC PT and metastatic sites (peritoneum and liver) suggested that distinct transcriptional programs are adopted by GCPM compared to liver mets. 333 differentially expressed genes found to discriminate between normal peritoneal and liver tissue^21^ were excluded to avoid confounding by admixture of resident tissue during sampling [**Supplementary Fig. 4c**]. Likewise, in view of the higher propensity for GCs of the TCGA CIN subtype to metastasize to the liver^14^, 1,172 differentially expressed genes discriminating between CIN and GS TCGA subtypes were excluded from the analysis to avoid confounding by intrinsic biological properties [**Supplementary Fig 4d**]. We found that the EMT signature and stromal infiltration appears to be uniquely enriched in PM. In a similar fashion, higher expression of memory activated CD4+ T cells, IFN iCAFs, ECM myCAFs and TGF myCAFs were uniquely enriched in PM. Conversely, high levels of M2 macrophages and monocytes were similarly high across both peritoneal and liver metastasis [**Fig. 4c**]. These findings suggest that GCPM is mechanistically distinct from hematogenous metastasis such as to the liver – consistent with previous findings in colorectal cancer (CRC).^4^

As tumors evolve towards PM, samples become increasingly mesenchymal^15^ (Fisher’s, p<0.001) and resembled a GI-mixed tumor lineage (Fisher’s, p=0.003) previously described by Wang *et al*^10^. The majority of PM samples originated from primary tumors with a Diffuse/NOS Lauren’s classification (Fisher’s, p<0.001). Tumor microenvironments^22^ were found to become increasingly fibrotic and less deserted in PM (Fisher’s, p=0.004) [**Fig. 4d**]. We also sought to evaluate the extent to which intrinsic genomic aberrations contribute to the process of transcoelomic metastases in GC. WES of 38 GCPMs, 51 PT with PM and 138 PT without PM were included. We found a significantly higher proportion of GS tumors in PM (Fisher’s, p<0.001), and lower proportion of samples with WGD (Fisher’s, p=0.005) and CNV (ANOVA, p<0.001). We also appreciated a non-significant trend towards increased monoclonality in PM samples (Fisher’s, p=0.119) [**Fig 4e**]. Analyses of driver mutations found a higher proportion of *CDH1* (Fisher’s, p=0.006), *POLE* (Fisher’s, p=0.091), *PIGR* (Fisher’s, p=0.113) mutations in PM samples. Conversely, *TP53*, *PIK3CA*, *KRAS* and *ERBB2* mutations were comparatively less common in PM samples [**Fig. 4f**].

### Putative targets in gastric cancer peritoneal metastasis

We then analyzed individual genes of clinical and therapeutic interest between paired and unpaired PT (n=49) and PM (n=100). Transcriptomic expression of emerging targets in GC such as the *CLDN18.2* isoform (paired T-test, p<0.001) and *FGFR2* (paired T-test, p=0.004) were found to be downregulated in GCPM. Conversely, we found upregulation of *NTRK2/NTRK3*, *PIK3CA*, *TEAD1/TEAD2, DKK1*^23^*, FGFR1 and HPSE* in the PM [**Fig. 5a, b**]. *HAVCR2 (TIM3)*, the co-inhibitory receptor expressed on IFN-γ-producing T cells and FoxP3+ Treg cells was found to be significantly upregulated in PM. No significant differences in gene expression programs relating other clinically relevant immune checkpoints of interest were found – *PDCD1 (or PD-1)*, *CD274 (or PD-L1)*, *CTLA4* and *TIGIT* [**Fig. 5a, b**].

**Fig. 5.**
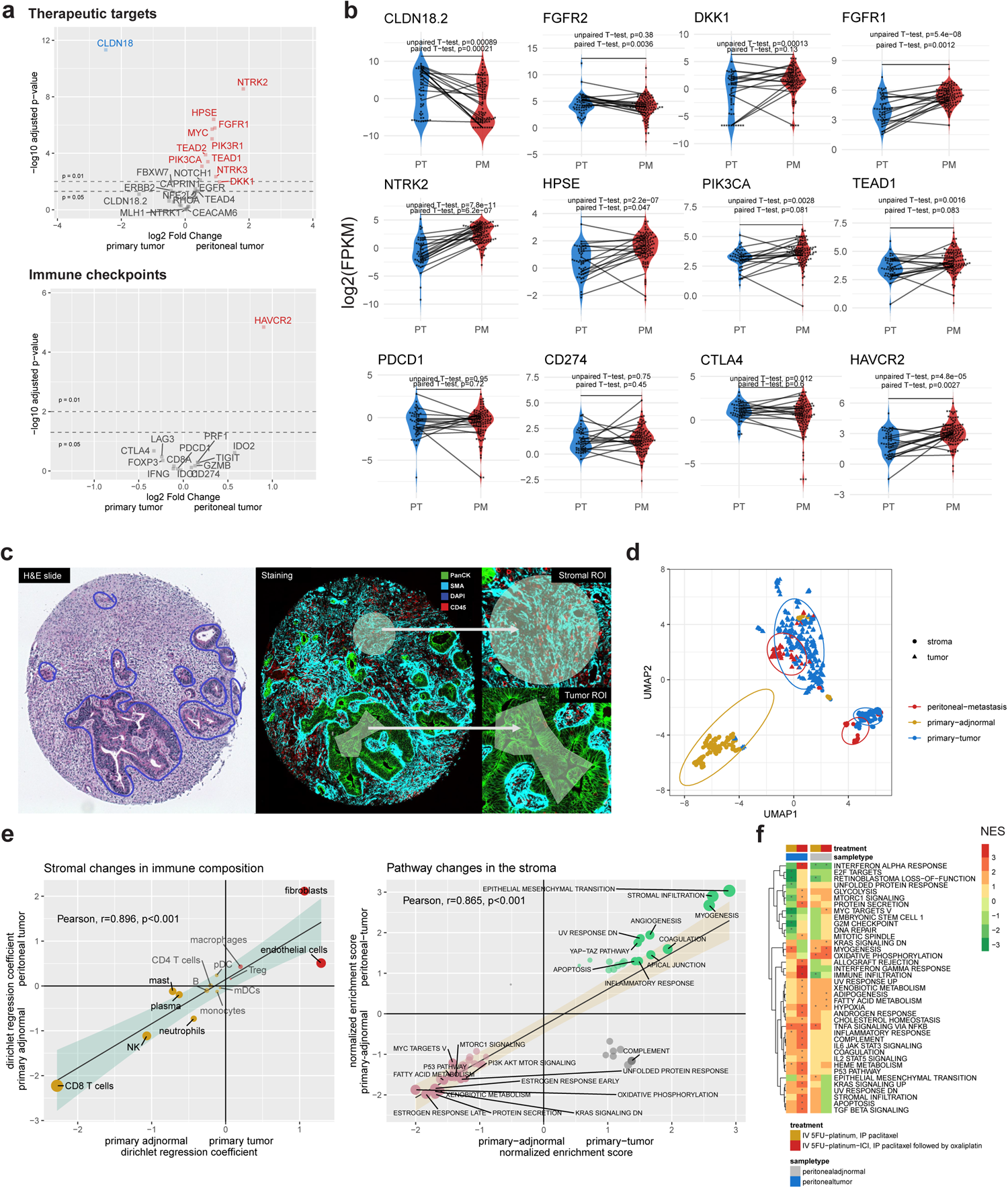
Putative targets and spatial transcriptomics of gastric cancer peritoneal metastasis. **a.** Volcano plot of putative targets between gastric cancer primary tumors and PM. The full list of included genes are provided in **Supplementary Table 6**. **b.** Beeswarm-violin plots of transcriptomic gene expression changes between primary tumor samples with PM (n=49) and PM samples (n=100). **c.** H&E slide, staining and selection of ROIs for DSP. **d.** UMAP of DSP data of gastric cancer samples, stratified by site of tumor and regions of interest (tumor vs stromal compartment). **e**. Scatter plots of Dirichlet regression coefficients of nuclei based immune cell proportions and GSEA comparisons from PT stroma vs primary adjacent normal stroma and PM stroma vs primary adjacent normal stroma comparisons. Cell types were highlighted if Bonferroni adjusted P-values were significant (p<0.05) in either comparison. Pathways were highlighted if Bonferroni adjusted P-values were significant in either comparison. **f.** Heatmap of pathway changes against treatment naïve samples. Values refer to GSEA NES for pathway comparisons and t-statistics from the unpaired t-test for immune cell type comparisons respectively. An asterisk refers denotes adjusted p-value <0.05. Abbreviations: PM, peritoneal metastasis; n=, number of samples; GSEA, Gene Set Enrichment Analysis; adjnormal, adjacent normal; NES, normalized enrichment score; IV intra-venous; 5FU, 5-fluorouracil; ICI, immune checkpoint inhibition; IP, intra-peritoneal.

### Spatial profiling demonstrates niche reprogramming within tumor and stromal compartments

Next, to understand the transcriptional dynamics of PT and PM tumor stromal compartments within their spatial context, we performed NanoString GeoMx DSP of tumor and stromal compartments of gastric PT and PM. Slides were stained for PanCK (epithelial cells), CD45 (immune cells) and SMA (stroma), and regions of interest (ROIs) were selected based on these markers [**Fig. 5c**]. A total of 712 ROIs were retrieved after quality control evaluation [**Supplementary Fig. 5a-c**; **Supplementary Table 5a, b**]. Unsupervised clustering of the ROIs revealed a clear distinction between adjacent normal and tumor, and between the tumor and stroma compartments. UMAP analysis demonstrated close clustering and overlap between the PT and PM for tumor ROIs, suggesting transcriptomic similarity [**Fig. 5d**].

Inspection of volcano plots between PT-PM tumor compartments did not identify any differentially expressed genes (log2[fold change]>2 and adjusted P-value<0.05) or major changes differences in *SpatialDecon* immune cell types [**Supplementary Fig. 5d, e**]. Transcriptomic similarity was likewise seen between PT-PM stromal compartments which appear distinct from adjacent normal primary stroma. [**Supplementary Fig. 5f, g**].

To understand the immune composition within the stromal compartments of the PT and PM, ROIs from these were compared against adjacent normal gastric stroma ROIs. Immune cell changes in both comparisons were shown to be concordant (Pearson’s R=0.896, p<0.001). *SpatialDecon* enumerated nuclei-based cell proportions of fibroblasts and endothelial cells were consistently higher in both cancer stromal compartments. Exploratory analyses of CAF subtypes^24^ (inflammatory CAF [iCAF], myofibroblast-like CAF [myCAF]) demonstrated higher expression of iCAFs compared to myCAFs in tumor stroma but not in primary adjacent normal stroma [**Supplementary Fig. 5h**]. Conversely, nuclei-based cell proportions of CD8 T cells were found to be lower in the cancer stromal compartment in both tumor types. Comparisons of pathway changes were also evaluated with GSEA. Consistent with the bulk RNA-seq analyses, EMT and stromal infiltration signatures were upregulated in the PT and PM stroma when compared to adjacent normal stroma. Pathway changes were concordant between both comparisons (Pearson, r=0.865, p<0.001) [**Fig. 5e**].

Overall, these findings suggest two possibilities. First, the PT could be undergoing modification of the microenvironment at the primary site, along with niche conditioning for transcoelomic seeding to occur. Second, the PT stroma could have developed pro-metastatic programs which were subsequently transferred into the peritoneum during metastasis. Unfortunately, it was not possible to confirm the first hypothesis due to the clinical nature of our datasets. Evaluation of “normal” peritoneal samples prior to the event of PM would be required and are not readily available through current clinical workflows.

### Effect of systemic and intra-peritoneal therapeutics on peritoneal samples

We investigated transcriptomic differences between GCPMs with paired grossly uninvolved peritoneum in GCPM retrieved prior to and after systemic and loco-regional intra-peritoneal therapeutics (PM n=58 samples; adjacent normal peritoneal samples in GCPM, n=36 samples). Patients were broadly recruited from early phase clinical trials that investigated intra-peritoneal paclitaxel or combination of PIPAC oxaliplatin/intraperitoneal paclitaxel with immunotherapy.^25,26^ No major shifts in transcriptional programs were detected in grossly uninvolved peritoneal tissue after exposure to systemic + IP therapeutics. However, peritoneal tumors treated with systemic chemotherapy + immune checkpoint inhibition (ICI) and IP chemotherapy demonstrated visible changes in examination of broad gene expression profiles through UMAP analysis [**Supplementary Fig. 5i**]. A deeper dive was performed on these gene expression programs using GSEA and immune deconvolution. No major pathway changes were detected between peritoneal-tumor samples after treatment with combination systemic and IP chemotherapy in contrast to treatment naive samples. However, in samples that were treated with systemic immunotherapy in addition to IP chemotherapy, we found a significant increase immune infiltration, inflammatory pathways (interferon gamma response, IL6-JAK-STAT, IL2-STAT5 signalling) and adipogenesis [**Fig. 5f**]. More M2 macrophages and monocytes were found in peritoneal tumor samples treated with systemic ICI [**Supplementary Fig. 5j**]. These findings highlight the importance of the synergy between combining locoregional and systemic therapies, particularly novel strategies that specifically target the tumor microenvironment.

### Niche reprogramming of the peritoneal environment in gastric cancer peritoneal metastasis

To better understand how the peritoneal environment might evolve to facilitate transcoelomic metastasis and peritoneal progression, we compared 53 adjacent normal peritoneal samples (grossly uninvolved at the point of sampling) in GCPM against 11 normal peritoneal samples from patients undergoing abdominal surgeries for various benign conditions. We found that adjacent normal peritoneum in GCPM was transcriptomically different from normal peritoneal samples [**Fig. 6a**] and was characterized by significantly increased myeloid dendritic cells, effector memory CD8+ T cells, CAFs and hematopoietic stem cells compared to normal peritoneum in benign conditions. There was also a trend towards increased M2 macrophages and Tregs [**Fig. 6b**]. Further exploration of CAF and dendritic cell subtypes show that majority of subtypes were enriched in the adjacent normal peritoneum in GCPM. Of note, ECM myCAFs (r=0.403, p=0.018) and plasmacytoid dendritic cells (r=0.304, p=0.080) were found to be correlated with higher PCI scores [**Fig. 6d**]. Our findings suggest that proliferation of these cell types within adjacent normal peritoneum are likely mechanistically involved in the progression of GCPM.

**Fig. 6.**
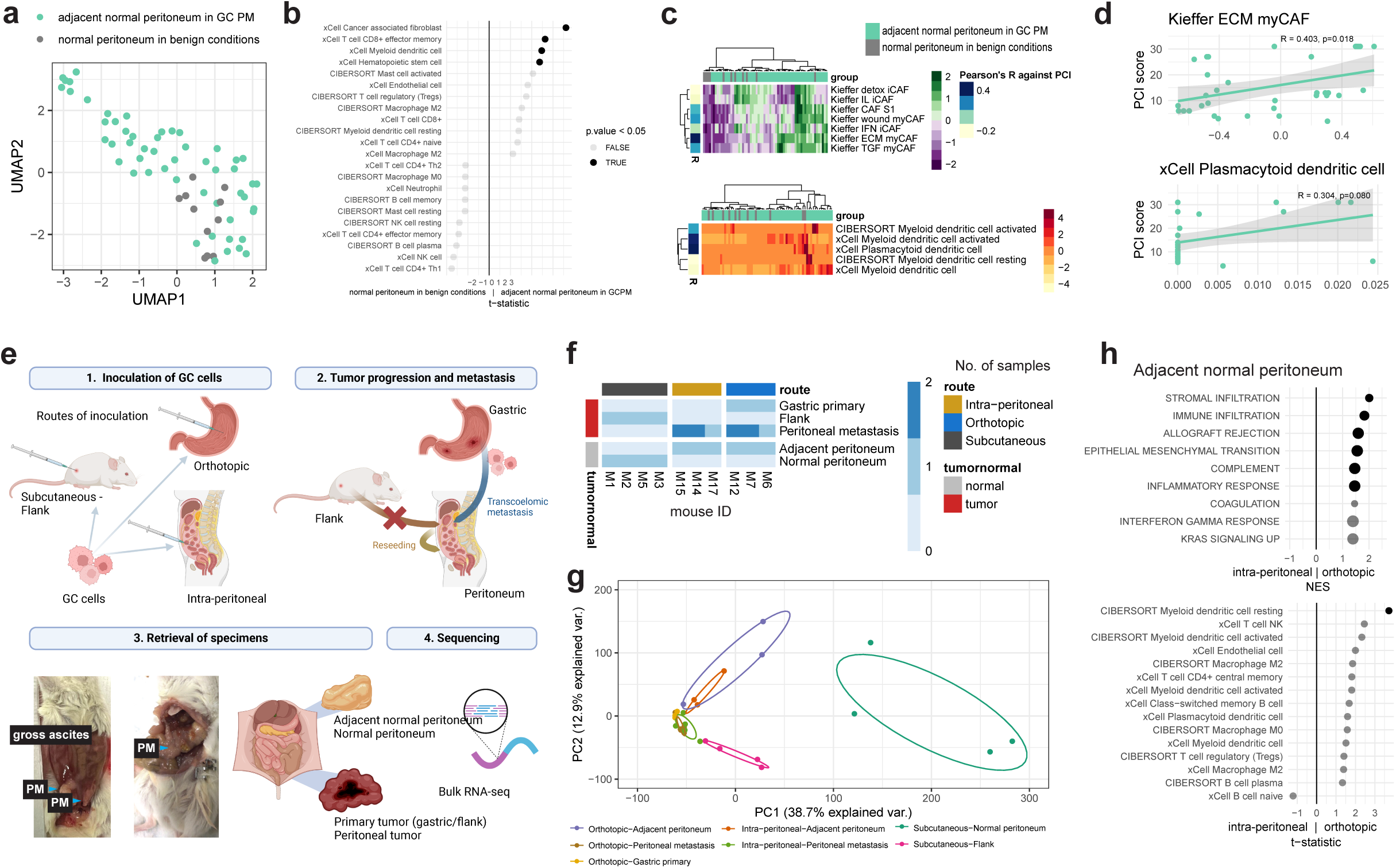
Niche reprogramming in gastric cancer peritoneal metastasis. **a.** UMAP of grossly uninvolved peritoneum in GCPM and normal peritoneum retrieved from patients with benign conditions. **b.** Comparisons of CIBERSORT/xCell enumerated cell types between grossly uninvolved peritoneum in GCPM and normal peritoneum in benign conditions. P-values were retrieved from an unpaired t-test. **c.** Heatmap of GSVA enrichment scores of CAF and DC subtypes. Correlation against PCI scores were conducted with Pearson’s correlation. **d.** Correlation of ECM myCAFs and plasmacytoid dendritic cell enrichment scores of grossly uninvolved peritoneal samples in GCPM against PCI scores. **e.** Overview of humanized mouse experimental design and workflow. The illustration was created using BioRender.com. **f.** Overview of samples collected from the 11 humanized mouse mouse models (n=4 flank innoculation; n=3 intra-peritoneal inoculation; n=3 orthotopic inoculation of GC cells) included in the study. **g.** PCA of samples from humanized mouse experiments. **h**. Dot plot of CIBERSORT/xCell cell type and pathway GSEA analysis between adjacent peritoneum from the intra-peritoneal and orthotopic humanized mouse model. **Abbreviations:** GC, gastric cancer; PM, peritoneal metastasis; PCI, peritoneal carcinomatosis index; ES enrichment score; myCAF, myofibroblastic-like cancer associated fibroblasts; NES, normalized enrichment score; ECM, extracellular matrix; GSEA, gene set enrichment analysis.

### Humanized mouse GC model suggests niche conditioning of the peritoneum in transcoelomic metastasis

We postulated that the peritoneum might undergo alterations during transcoelomic metastasis to facilitate homing of tumor seeds. This concept, first introduced by Stephen Paget in 1889 as pre-metastatic niche conditioning, remains unconfirmed in gastrointestinal (GI) peritoneal metastases^27^. To address this gap, we utilized an innovative humanized mouse model as the host immune system appears to play a significant role in the mechanisms of transcoelomic metastases.^9^ Humanized mice (humice) were injected with GSU (diffuse type) GC cell lines in the flank (subcutaneous) (n=4), stomach (orthotopic) (n=3) and directly into the peritoneum (n=3). After approximately 1 month, humice were sacrificed and samples from the PT, PM and adjacent peritoneum were retrieved [**Fig. 6e**]. Consistent with previous studies demonstrating higher propensity for orthotopic transplantation for metastases compared to the subcutaneous route^28–31^, all 3 orthotopic humice were found to have PM while no gross PM were appreciated in the flank humice model. Peritoneal samples from these flank humice were retrieved and labelled as “normal peritoneum” [**Fig. 6f**]. The tumors were then harvested, along with adjacent normal peritoneum and bulk RNA-seq was performed and analyzed [**Supplementary Fig. 6a, b**].

Principal component analysis (PCA) mapping samples demonstrated segregation between normal peritoneum from the flank model and adjacent normal samples in the orthotopic model and intra-peritoneal model. Tumor samples of PT and PM samples clustered closely and distinctly from PT and PM samples from the flank humanized mouse model, suggesting transcriptomic similarity as alluded to in our human tumor cohorts [**Fig. 6g**]. Compared to adjacent normal peritoneum of the intra-peritoneal humanized mouse model, the orthotopic model revealed enrichment of tumorigenic programs such as EMT, stroma and immune infiltration, and inflammatory response signatures [**Fig. 6h**]. Enrichment scores of these pathways (in addition to upstream identified signatures such as T cell exhaustion, angiogenesis and inflammatory pathways [IL2-STAT5, IL6-JAK-STAT3, TGF-β signaling]) were likewise more pronounced in adjacent normal peritoneal samples in the orthotopic humanized mouse model when contrasted against other sample types [**Supplementary Fig. 6c**]. We also found a trend towards higher myeloid dendric cells, endothelial cells, M2 macrophages, class-switched memory B cells and Tregs in adjacent peritoneum from the orthotopic model (n=3) when compared to the intra-peritoneal model (n=3) [**Fig. 6h, Supplementary Fig. 6d**]. Overall, findings from the humanized mouse experiment demonstrate concordance with our analyses of human samples, providing initial evidence of niche reconditioning of the peritoneum in transcoelomic metastasis.

## Discussion

The poor prognosis of GCPM is typically attributed to, and characterized by, resistance to systemic chemotherapy and immunotherapy.^32^ The evolutionarily conserved plasma-peritoneal barrier as well as reduced tumor tissue vascularity contributes to poorer responses to systemically administered antineoplastic therapy.^33^ To our knowledge, ours is one of the largest studies to date with GCPM tissue specimens (compared to malignant ascites), paired with PT, analyzed using WES, WTS and spatial profiling. Our study identified molecular markers and pathways predictive of GCPM, along with gene expression signatures associated with the tumor microenvironment (TME) such as stromal infiltration and M2 macrophages. We delineated distinct pathways and immune compositions in GCPM compared to liver metastases, underlining the significance of TME changes in transcoelomic metastases. Additionally, we found differential expression of therapeutic targets between primary tumors and PM, emphasizing the importance of combining locoregional and systemic therapies targeting the TME. Our study also revealed niche reprogramming in GCPM, indicating mechanistic involvement of myeloid dendritic cells, effector memory CD8+ T cells, and cancer-associated fibroblasts in metastatic progression. Utilizing a humanized mouse model, we provide initial evidence of niche conditioning of the peritoneum during transcoelomic metastasis.

Previous studies have described predictors of GCPM, however must of these have been limited to clinico-pathologic markers. These include characteristics such as T and N stage of tumor, histological subtype, tumor markers, and radiological features.^34–36^ Spatial transcriptomic analysis of GC support metastatic dissemination of cancer from the serosal (rather than mucosal) surface of the PT.^37^ However, identifying occult peritoneal metastases through imaging can be difficult and the current standard-of-care remains a diagnostic laparoscopy. Few cohorts have been developed to prospectively follow up patients for peritoneal outcomes. Our study identified *CDH1* mutations as a predictor of PM, which has been reported previously.^38^ However, our study also identified *PIGR* mutations as a predictor of GCPM. *PIGR* encodes the polymeric immunoglobulin receptor, a member of the immunoglobulin superfamily involved in transporting IgA proteins across mucosal epithelia. *PIGR* also functions as a precursor of the secretory component of IgA which regulates immune function.^39^ Interestingly, in non-malignant tissues associated with ulcerative colitis (UC), somatic *PIGR* mutations have been reported.^40^ Specifically, clones carrying *PIGR* mutations have been shown to be able to avert *IL17*-induced cytotoxicity and selectively expand in inflamed colonic environments, notably without leading to cancer. The presence of *PIGR* indel and non-sense mutations in UC samples is very similar to the mutation pattern observed in GC samples, suggesting a possible link between inflammation-related mutation and cancer progression.

Our interpretation of WES analyses aligns with the alterations observed in the TME through WTS analyses, revealing that gene expression signatures associated with the TME, including stromal infiltration, epithelial-mesenchymal transition, cancer-associated fibroblasts, and M2 macrophages, are correlated with a higher incidence of PM. The Hippo pathway was found to be associated with the transcoelomic development of PM from PT and upregulated in PM samples. The Hippo pathway controls multiple cellular functions that are central to tumorigenesis, including proliferation and apoptosis.^41^ Several therapeutic strategies are being pursued in targeting the Hippo pathway, including *TEAD1* and *YAP-TAZ*, with promising preclinical and preliminary data being developed.^11,42^ With multiple cross-linking pathways and interactions between Hippo, TGF-β, EMT and cancer associated fibroblasts, nuanced disruptions of these pathways lead to changes in the TME and form potential immunotherapeutic vulnerabilities.^43^

Our study also compared the differential expression of various therapeutically relevant genes and pathways between PT and PM as well as, for the first time, before and after exposure to intraperitoneal therapy. We also compared GCPM with liver metastases. We found several clinically relevant and important findings such as lower expression of *CLDN18.2* in PM compared to PT. This may have immediate impact, as *CLDN18.2* inhibitors are poised for regulatory approval.^44,45^ Considering the distinct biology of transcoelomic metastases, compared to hematogenous spread, as elucidated in our study, these findings form the grounds for several changes to be considered in the design of future clinical trials involving advanced or metastatic GC. First, clinical trials involving targeted therapies should examine variation in expression of the putative target in PM compared to PT, and if unsure or different, should consider including presence of PM as a stratification factor. Second, trials involving immunotherapies should consider the distinct immune niche of PM and use stratification factors, and/or develop trials that are PM specific. Third, the mechanisms of transcoelomic metastases elucidated in our study support the development of adjuvant strategies specific for the prevention of PM, which possibly require the administration of locoregional therapies directly into the peritoneum. The demonstration of the development of a premetastatic niche within the peritoneum supports the principles of early disruption of the communication between the PT and the metastatic site. Lastly, the distinct biological programs that occur in the PM, within the immune niche, described in our study support the development of peritoneal-specific novel therapeutic strategies including direct intraperitoneal administration of chimeric antigen receptor T-cell (CAR-T) therapies and oncolytic virotherapies.^46,47^

Our study has several limitations. To fully study niche conditioning, “normal peritoneum” from early GC patients that have not developed peritoneal metastases would be required. Acquiring these tissues would require new ethics approval, as sampling of the uninvolved peritoneum in early-stage patients is not performed in routine clinical practice. Further, longitudinal follow up is required to distinguish how normal peritoneum of patients with early GC defers between patients with and without eventual PM. As such, we were unable to answer the temporal question of pre-metastatic niche conditioning at this juncture. Additionally, experimental models need to be developed to definitively prove several of the genomic predictors of GCPM, with the current findings being associative rather than mechanistic. Lastly, single-cell sequencing of PM would have provided a more granular insight into the PM TME. While this was attempted, due to the fibrous nature of many of the PM samples, it was not technically feasible. We previously reported single-cell sequencing results from a limited number of PM samples, and other studies have reported findings from single-cell sequencing of malignant ascites.^48^

The identification of predictive markers and therapeutic targets in GCPM opens promising avenues for targeted interventions and improved patient outcomes. Understanding the role of the tumor microenvironment (TME) alterations, offers new opportunities for innovative treatments. Moving forward, integrating these insights into clinical trial designs and developing peritoneal-specific therapies hold the potential to transform the management of PM, offering hope for patients facing this challenging condition. While there are existing limitations to address and further research is needed for validation, these findings pave the way for a more hopeful future in combating GCPM.

## Methods

### Clinical Cohorts

#### Prospective gastric cancer primary tumor cohort

Patients with newly diagnosed GC at National University Hospital, Singapore and Tan Tock Seng Hospital, Singapore, were prospectively enrolled with informed consent (Gastric Cancer Biomarker Discovery II, GASCAD II, DSRB Ethics Approval No: 2005/00440 and NUS-IRB LH-19-070E). GC tissue samples were obtained with clinical and pathologic annotation from January 2006 to December 2015. Staging information was determined histopathologically and in combination with clinical information. American Joint Committee on Cancer (AJCC) (8th edition) on GC staging and Lauren’s classification of GC were used. Patients with less than 3 months of follow-up for recurrence or death were excluded. Samples were retrieved and quality checks performed prior to downstream processing. Patients were prospectively followed up for recurrence in the peritoneum and/or other organs, which was detected through clinical, radiological and/or diagnostic laparoscopy. All recurrences were discussed at a multidisciplinary tumor board for concordance.

#### Cross sectional cohort: gastric cancer

Patients with GCPM had PM and adjacent normal tissue sampled in the National University Hospital Singapore, and the National Cancer Center, Singapore. Patients were prospectively consented for tissue and clinical data collection on protocols: (Feasibility Study of Intraperitoneal Paclitaxel With Oxaliplatin and Capecitabine in Patients With Advanced Gastric Cancer (DSRB Ref: 2012/00429); Phase 1 Study of Pressurized Intraperitoneal Aerosol Chemotherapy (PIPAC) with oxaliplatin and in combination with Nivolumab in patients with peritoneal carcinomatosis (PIANO) (DSRB Ref: 2016/01088); Using QPOP to predict treatment for GastroIntestinal Cancers (Q-GAIN) (DSRB Ref: 2019/00924); Gastric Cancer Biomarker Discovery (GASCAD-II) (DSRB Ref: 2005/00440); Circulating Markers in Gastrointestinal and Hepatopancreatobiliary Cancers (CIRB 2018/3046); INtegrated Diagnostics and discovery In Gastro-Oesophageal Cancers (CIRB 2015/2363)).

#### Liver metastases cohort: Gastroesophageal Cancer

Retrieval of samples and RNA-sequencing of samples from this cohort has been previously described by Damhofer *et al.*^20^. Bioinformatic pipelines of processing FASTQ files were harmonized with the other datasets in this study and downstream gene expression matrices were integrated into current analyses.

#### DNA extraction and whole exome sequencing

Genomic DNA from 20 mg of snap-frozen resected gastric tissue was extracted using the QIAamp DNA mini kit (Qiagen, USA). DNA extraction was performed according to the manufacturer’s protocol. Purified genomic DNA was stored at −20°C. The Agilent SureSelect Human All Exon v6 kit (Agilent Technologies) was used for whole exome sequencing. Using 300 ng of DNA, whole exome-sequencing was performed on the Illumina HiSeq platform to generate 150 bp paired-end sequencing reads (Novogene-AIT, Singapore).

#### Mutation calling and mutational signatures

Exome sequencing reads were aligned to the reference human genome hs37d5 using bwa mem^49^. Preprocessing steps, including duplicate marking, local read realignment and base quality score recalibration were performed using Picard and Genome Analysis Toolkit (GATK)^50^ to generate analysis-ready BAM files. Mutect2^51^ was used in paired mode to generate somatic SNVs and indels by comparing BAM files from tumor and matched normal or blood samples. Germline variants were filtered using the gnomAD database and a Panel of Normals generated from all normal samples. Sequence context artifacts were filtered using the *FilterMutectCalls* module. Mutation calling in GC driver genes were also performed using Strelka^52^ to identify somatic mutations missed in Mutect2. Somatic mutations were annotated using Funcotator. For mutational signature analysis, we fitted GC-specific mutational signature to each samples using the *signature.tools.lib* package ^53^ in R. Analysis of somatic variants were conducted with the *maftools* package.^54^ Comparisons of subgroups to identify mutations associated with PM were undertaken with the *mafCompare* function. Somatic interactions were evaluated with the *somaticInteractions* function.

#### Classification of TCGA subtypes

GCs were first classified as EBV or MSI. The remaining tumors were then classified as either CIN or GS based on the presence of whole-genome doubling. Samples were classified as EBV by evaluating the number of reads aligning to the NC_007605 EBV genome (included in the hs37d5). MSI status was assessed using MSIsensor2^55^. Samples with MSI scores of more than 10 were classified as MSI positive. We used GISTIC2^56^ to identify significant somatic copy number alterations in GC samples. For assignment of CIN/GS subtypes, tumors were clustered using Euclidean distance and Ward’s method, based on thresholded copy number from significantly altered sites identified from GISTIC2 (q<0.05). WGD calls were performed using Facets^57^. Samples with major copy number fraction across the autosomal genome that is greater than 0.5 were considered as WGD^+^. GC samples in cluster enriched with WGD^+^ were assigned as CIN.

#### RNA extraction and whole transcriptome sequencing

For RNA-seq experiments, total RNA was extracted using the RNeasy Mini Kit (Qiagen), and library preparation was conducted using the Tru-Seq Stranded Total RNA with Ribo-Zero Gold kit protocol (Illumina). Libraries were sequenced on a HiSeq4000 sequencer using the paired-end 150 bp read option. QC-passed reads were aligned to the human reference CGRh38/hg38 genome using STAR v.2.7.9a. Transcript abundance quantification was performed using RSEM v1.3.3.^58^ RNA-seq data were normalized by Log2 FPKM unless stated otherwise.

#### Gene expression analysis of RNA-seq data

##### Batch effect adjustments and evaluation

Batch correction after batch integration was conducted with ComBat-Seq^59^ with covariants including sample site (primary vs peritoneum) and sample type (normal vs tumor). Post batch correction evaluation was conducted with principle component analysis of housekeeping genes.^60^

##### RNA-seq analyses

Differential analysis of count data was conducted with the *DESeq2*^61^ package. Tumor purity of tumor samples was evaluated with the ESTIMATE algorithm.^62^ Dimension reduction of RNA-seq and DSP data was conducted with Uniform Manifold Approximation and Projection (UMAP). Ellipses were added to UMAP plots for visualisation of cluster overlap and/or segregation with the *ggbiplot* package. Trajectory and pseudotime analyses were primarily utilized for visualization and were undertaken with the *slingshot*^63^ package in R. Each trajectory was anchored by clusters found to have the greatest proportion samples originating from the earliest parental tumor (i.e. GC: PT without PM).

Pathway enrichment scores were retrieved with single sample GSEA^64^ or GSVA utilizing the GSVA package^65^ in R-4.2.0. The Molecular Signatures Database (MSigDB) hallmark^66^ gene signature set was utilized for the primary analysis, additional gene signature sets utilized are documented in **Supplementary Table 6**. Immune cell subsets were enumerated primarily with the CIBERSORT^67^ LM22 immune subset signature and xCell^68^. These were implemented through the *immunedeconv* package^69^. Further exploratory analyses of immune cell types not available within these packages were conducted with ssGSEA enrichment scores of previously reported gene signatures [**Supplementary Table 6**]. Tumor microenvironment subtypes were retrieved by methods previously described by Bagaev *et al.*^22^, with the pan TCGA cohort utilized as the training model, analyses were conducted in python.

To identify CIBERSORT/xCell enumerated immune cell types and pathways involved in PM, correlation analysis with Spearman’s test was conducted across PT without PM, PT with PM and PM samples.

#### Spatial transcriptomic analysis

Specimens from patients with GC were prepared as either whole slides or as a tissue microarray (TMA). Primary and peritoneal FFPE tissues were mounted on Leica Bond slides for the NanoString GeoMx DSP platform. For H&E staining, FFPE slides were deparaffinized with histoclear, rehydrated and stained with Hematoxylin Solution. Slides were counterstained with eosin and mounted with mounting media. FFPE slides were subjected to conventional tissue preprocessing (deparaffinization and rehydration). Standard fluorescence-labeled morphology marker panel consisting of Pan-CK for epithelial regions, CD45 for immune cells, α-smooth muscle actin for fibroblast and nuclear stain were used as ROI selection references. ROIs for each slide were drawn and selected.

Data were analyzed by uploading the counts data set from the Illumina run into the GeoMx DSP analysis suite. Biological probe QC was performed using default settings. Quality control and pre-processing of digital spatial profiled next generation sequenced data (DSP-NGS) was undertaken with packages *NanoStringNCTools*, *GeomxTools* and *GeoMxWorkflows*. Samples with raw sequencing reads with >1000 raw reads, <80% alignment, trimmed or stitched reads, no template control (NTC) counts greater than 10,000, <50% saturation and <150 nuclei per segment were removed. Batch correction was undertaken with *CombatSeq*. Q3 averaged normalized DSP-NGS GeoMx mRNA expression was utilized for downstream analyses. Cell abundance (nuclei derived count) and proportion estimates were retrieved with the *SpatialDecon*^70^ package. The default safeTME cell profile matrix was referenced. For analyses of cell type ratios, continuity correction with a proportion of 5e-05 was implemented. Dirichlet regression of cell proportions was utilized as the primary approach to elucidate differences between sample types.

#### Humice experiments

##### Humanised mice model

All experiments and procedures pertaining to animals were approved by the Institutional Animal Care and Use Committee (IACUC #191440) of the Agency for Science, Technology and Research (A*STAR), Singapore. Generation of mice for experimentshas been previously described.^71^ In brief, 1-3 day old NOD-scid Il2rγ^null^ (NSG) pups were irradiated with sublethal dose (1 Gy) and thereafter engrafted with 1×10^5^ human CD34^+^ cord blood cells (HLA-A24:02, Stemcell Technologies) through intrahepatic injection. Mice were evaluated 10 weeks after engraftment and submandibularly bled to assess the reconstitution of human immune system (humanised) by flow cytometry. Only mice which had >10% humanised (proportion of human CD45 cells: total CD45 cells) were deemed suitable for this experiment.

##### Experimental design

To evaluate the varying gene expression programmes required to establish metastases at differing sites, a diffuse-type cell line of HLA-A24:02 subtype (GSU) was inoculated into bilateral flanks, via intra-gastric (orthotopic) injection and direct intraperitoneal (IP) injection of humanised mice. Orthotopic inoculation of GC cell lines has been shown to produce PM that recapitulate the original PM.^72^ GSU cells were prepared and standardised to 30ul with 50% Matrigel diluted in saline. Five million GSU cells were injected into the subcutaneous plane of each flank of the mice. In orthotopic and IP inoculation, 2×10^6^ cells were introduced either into the gastric wall or directly into the peritoneal cavity. To investigate differences in the tumor microenvironment, the above experiment was repeated in NSG mice as well. Mice had *ad libitum* access to food and water, were evaluated weakly and sacrificed approximately 1 month after inoculation. A careful necropsy was performed and relevant tumors from the flank, stomach and peritoneum were harvested. In all experimental animals, the peritoneum was carefully inspected for gross lesions. In animal subjects with PM, the adjacent peritoneum was also harvested for further evaluation.

#### Statistics and reproducibility

A two-sided T-test was used for comparisons of continuous variables between two groups with normalized distributions, and a two-sided Wilcoxon test was used when variables were not normally distributed. Comparisons between more than two groups were performed by analysis of variance (ANOVA). For comparisons of count variables, the Fisher’s exact test was utilized for unpaired samples while the McNemar’s test was utilized for paired samples. Comparisons of proportions was undertaken with a univariate Dirichlet regression model with the package *DirichletReg* in R. The strength of correlation was measured using the Pearson (P) or Spearman (Rho, ρ) correlation coefficient and the probability of observing a correlation with the corresponding P-values. Survival analyses for prediction of peritoneal metastases were shown with Kaplan-Meier plots and conducted with a Cox-proportional hazards model with Fine-Gray model to account for competing risks of death using the *survival* package. The log-rank test was used to derive P-values for survival analyses. Exact P-values were provided whenever possible. P-value adjustment was conducted with the Bonferroni correction unless otherwise stated.^73^ No statistical method was used to predetermine the sample size for this study. Sample size was largely limited by the size of the samples provided and successfully assayed for this study.

All analyses were conducted in R-4.2.0 unless stated otherwise. Graphical illustrations were created with BioRender.com. Supplementary Information is available for this paper.

## Author contributions

**J. J. Zhao:** Conceptualization, software, formal analysis, investigation, project administration, visualization, methodology, validation, writing–original draft, writing–review and editing.

**J. C. A. Ong**: Data curation, funding acquisition.

**S. Srivatsava**: Data curation.

**D. K. A. Chia**: Methodology, data curation, writing–review and editing.

**H. Ma**: Investigation, formal analysis.

**K. Huang**: Investigation, formal analysis.

**T. Sheng**: Investigation, formal analysis.

**K. Ramnarayanan**: Data curation.

**X. Ong**: Data curation.

**S. T. Tay**: Data curation.

**T. Hagihara**: Data curation.

**A. L. K. Tan**: Data curation.

**M. C. C. Teo**: Data curation.

**Q. Xuan**: Data curation.

**G. Ng**: Data curation.

**J. Tan**: Data curation.

**M. C. H. Ng**: Data curation.

**A. Shabbir**: Data curation.

**G. Kim**: Data curation.

**Y. Tay**: Funding acquisition, conceptualization.

**Z. Her**: Data curation.

**G. Leoncini**: Data curation.

**B. T. Teh**: Data curation.

**J. H. Hong**: Data curation.

**R. Tay**: Formal analysis.

**C. B. Teo**: Formal analysis.

**M. P. G. Dings**: Data curation, writing–review and editing.

**M. Bijlsma**: Data curation, writing–review and editing.

**Y. X. Gwee**: Data curation.

**R. Walsh**: Data curation.

**J. H. Y. Lum**: Data curation.

**J. H. Law**: Data curation.

**F. Pietrantonio**: Data curation.

**S. M. Blum**: Writing–review and editing.

**H. V. Laarhoven**: Data curation, writing–review and editing.

**S. J. Klempner**: Writing–review and editing.

**W. P. Yong**: Conceptualization, data curation, writing–review and editing.

**J. B. Y. So**: Data curation, writing–review and editing.

**Q. Chen**: Data curation.

**P. Tan:** Conceptualization, supervision, funding acquisition, project administration, writing– review and editing.

**R. Sundar**: Conceptualization, data curation, supervision, funding acquisition, project administration, writing–original draft, writing–review and editing.

## Acknowledgements

The authors would like to thank Ms Hazel Hng from the National University of Singapore, Singapore for assisting with project administration for the conduct of this study.

