## Supplementary Figures for "Spatially resolved niche and tumor microenvironmental alterations in gastric cancer peritoneal metastases"

**SUPPLMENTARY FIGURES**

**CONTENT PAGE**

[**Supplementary Figure 4c** Volcano plot of differential gene expressions between GTEx normal peritoneal (n-541) and liver (n=226) samples– 333 differentially expressed genes (adjusted p-value <0.05 & |log2FC| >2) were identified.	8](#_Toc161005839)

[**Supplementary Figure 4d** Volcano plot of differential gene expressions between CIN (n=99) and GS (n=87) primary gastric cancer samples from the prospective cohort – 1172 differentially expressed genes (adjusted p-value <0.05 & |log2FC| >2) were identified. Samples were retrieved from the TCGA database.	8](#_Toc161005840)

Remarks: Figures were numbers according to the relevant figures it was referenced to in the main manuscript. For example, the content of **Supplementary** **Figure 2** is relevant to main **Figure 2**.

### SUPPLEMENTARY FIGURES

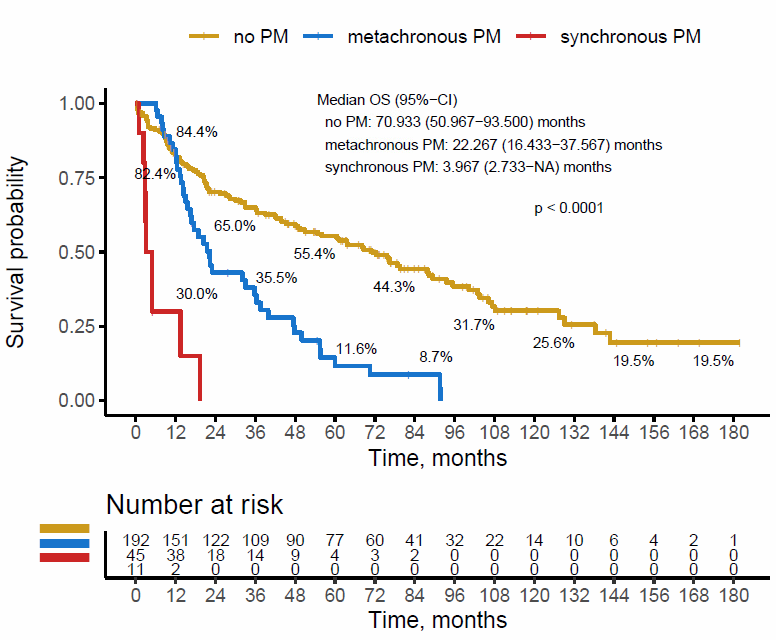

#### **Supplementary Figure 2a** Overall survival of the prospective cohort stratified by onset of peritoneal metastasis.

Remarks: P-values were retrieved from the log-rank test. t=0 is defined as the time of diagnosis. Synchronous PM is defined as the diagnosis of PM within 6 months of diagnosis of gastric cancer, while metachronous PM is defined as the diagnosis of PM after 6 months of diagnosis of gastric cancer. Median follow up time was 87.0 months.

Abbreviations: OS, overall survival; CI, confidence interval; PM, peritoneal metastasis

**b**

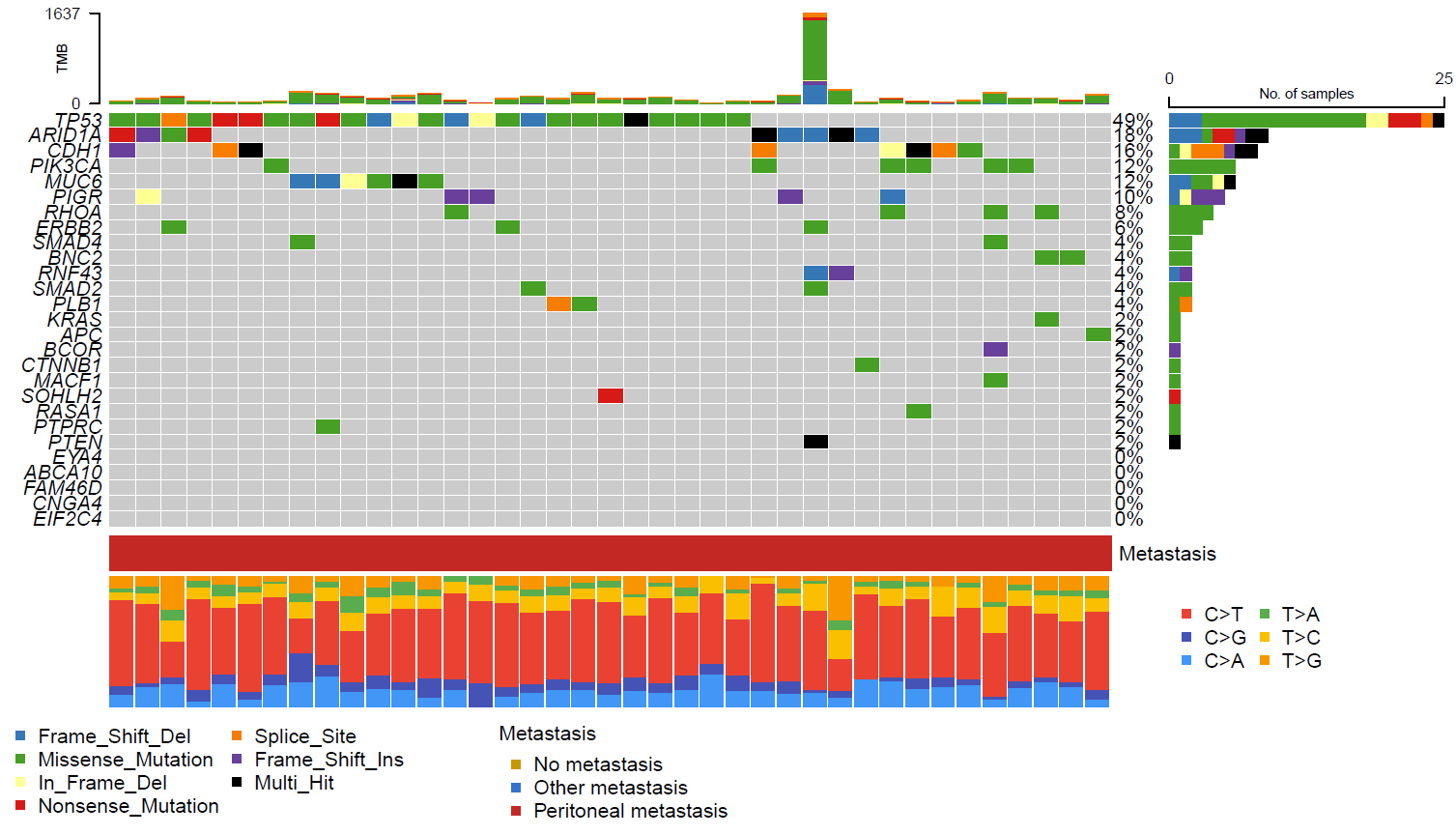

**c**

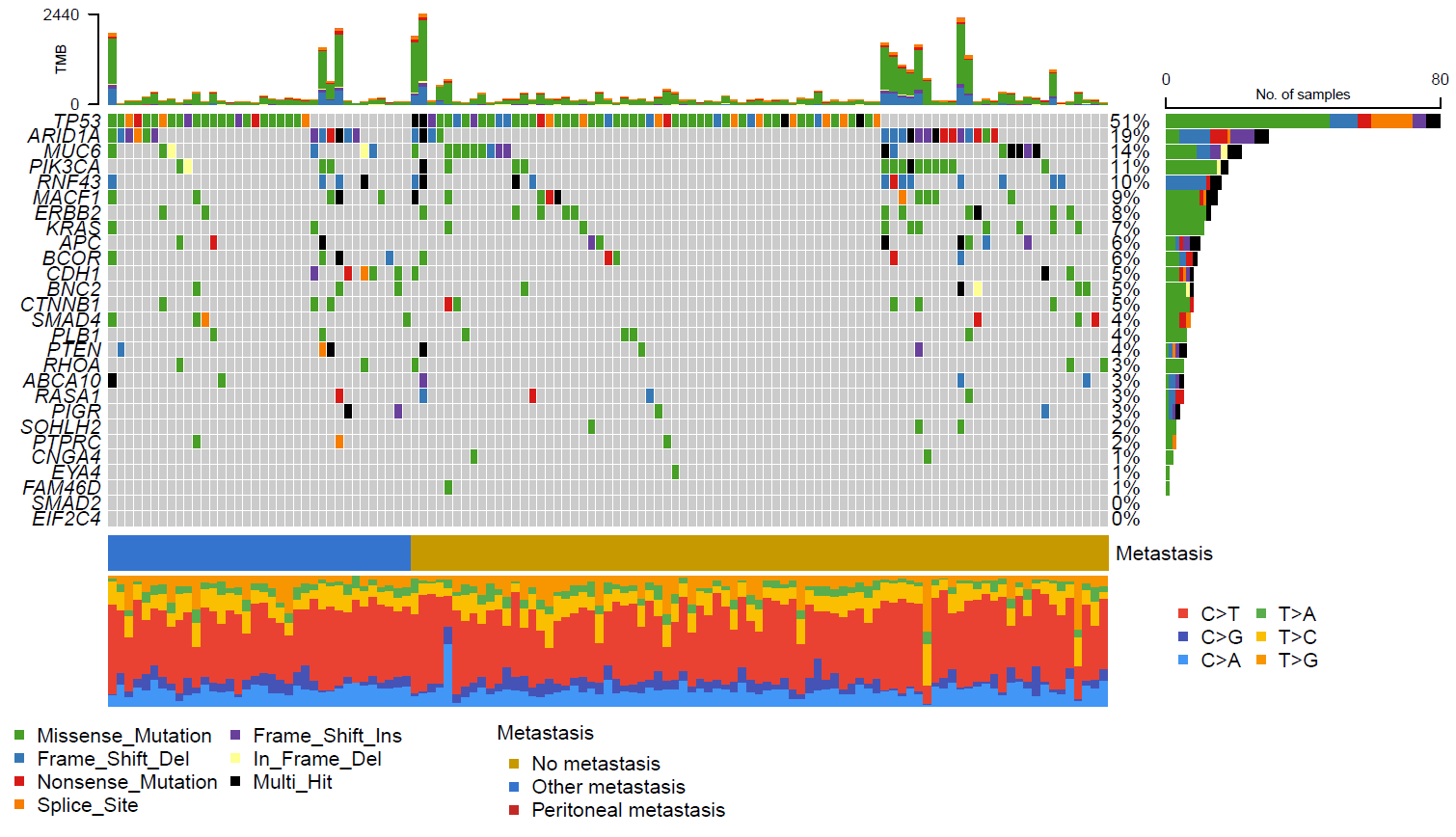

#### **Supplementary Figure 2b, c** Oncoplot of top 30 genetic alterations found in patients with (b) peritoneal metastasis and (c) without peritoneal metastasis in the prospective cohort

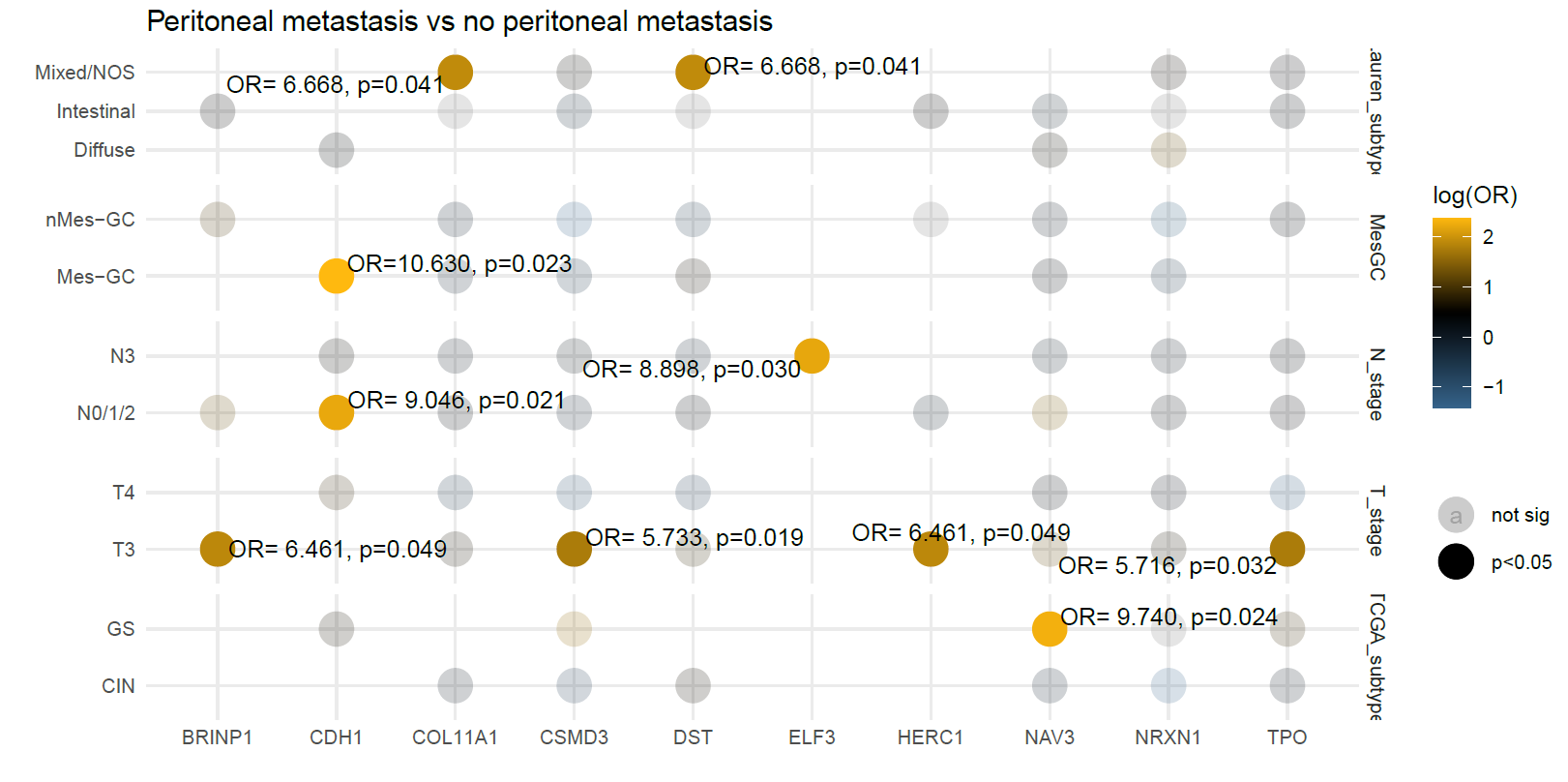

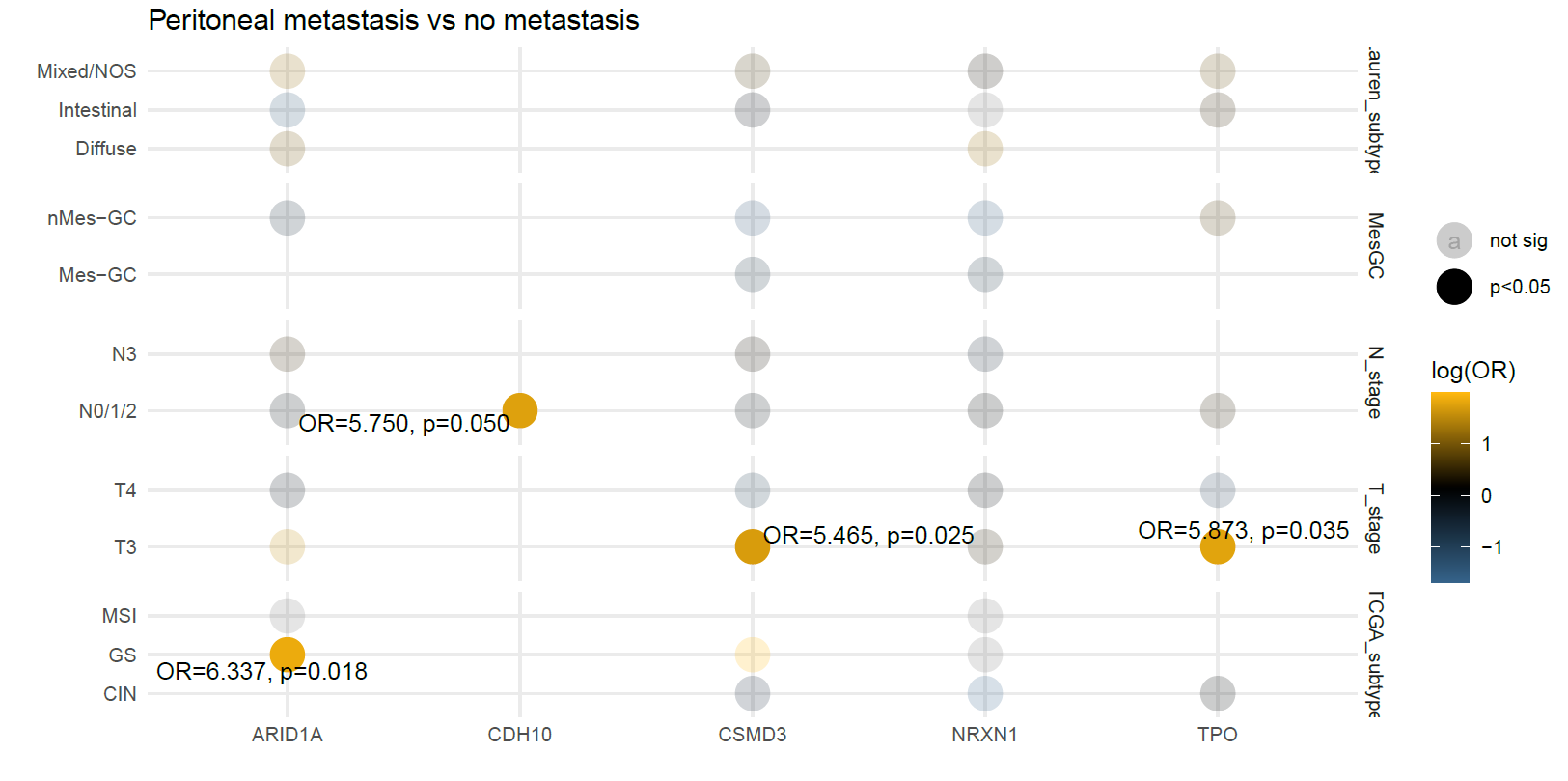

#### **Supplementary Figure 2d** Subtype analysis altered genes (all genes included) of primary tumours in gastric cancer predictive of peritoneal metastasis

Abbreviations: OR, odds ratio; sig, significant; TCGA, The Cancer Genome Atlas; GC, gastric cancer; Mes, mesenchymal subtype; nMes, non-mesenchymal subtype; NOS, none otherwise specified; CIN, chromosomal instability; GS genomically stable; MSI, microsatellite instability; EBV, Epstein-Barr virus

| **a** | **b** |
| --- | --- |
| 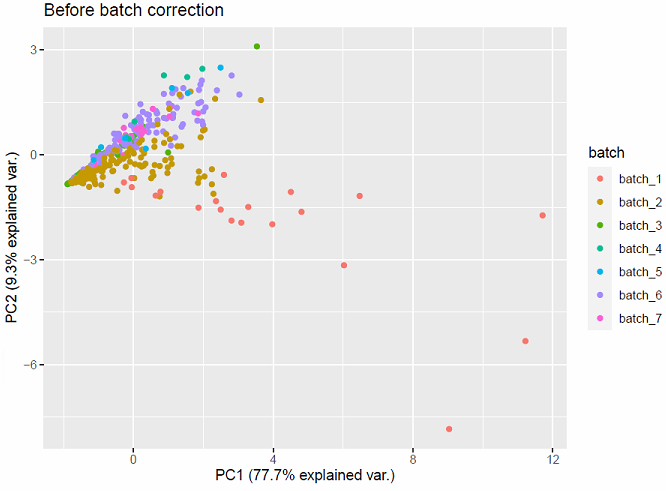 | 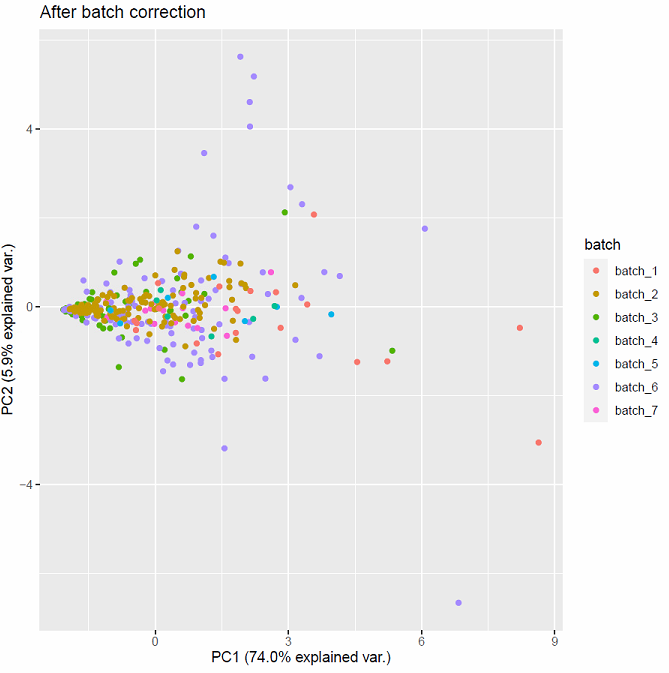 |

#### **Supplementary Figure 4a, b** Batch integration and batch correction evaluation of integrated WTS data of cohorts

Abbreviations: PCA, principle component analysis; var., variance; WTS, whole transcriptomic sequencing.

| **c** |
| --- |
| 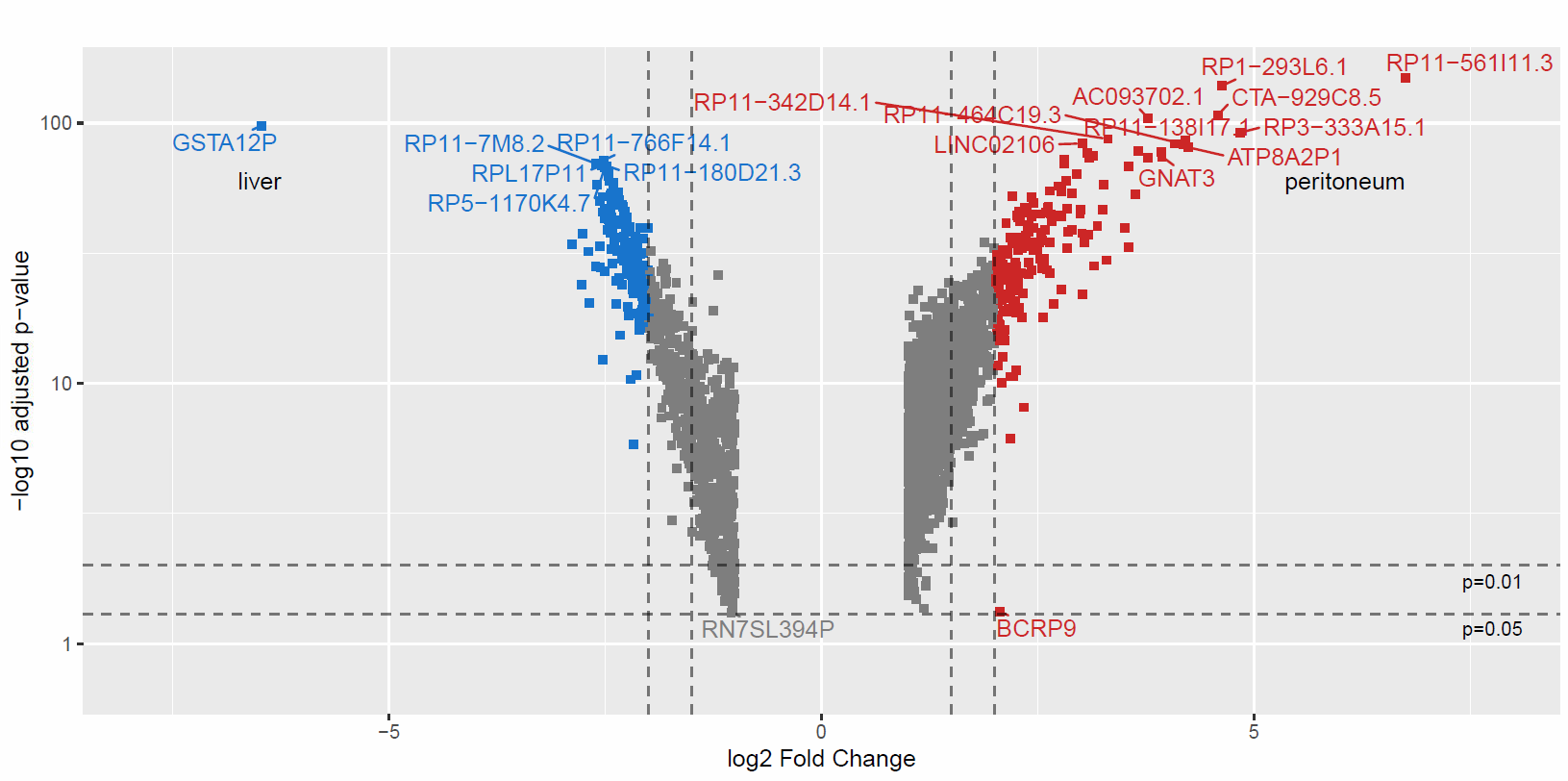 |
| **d** |
| 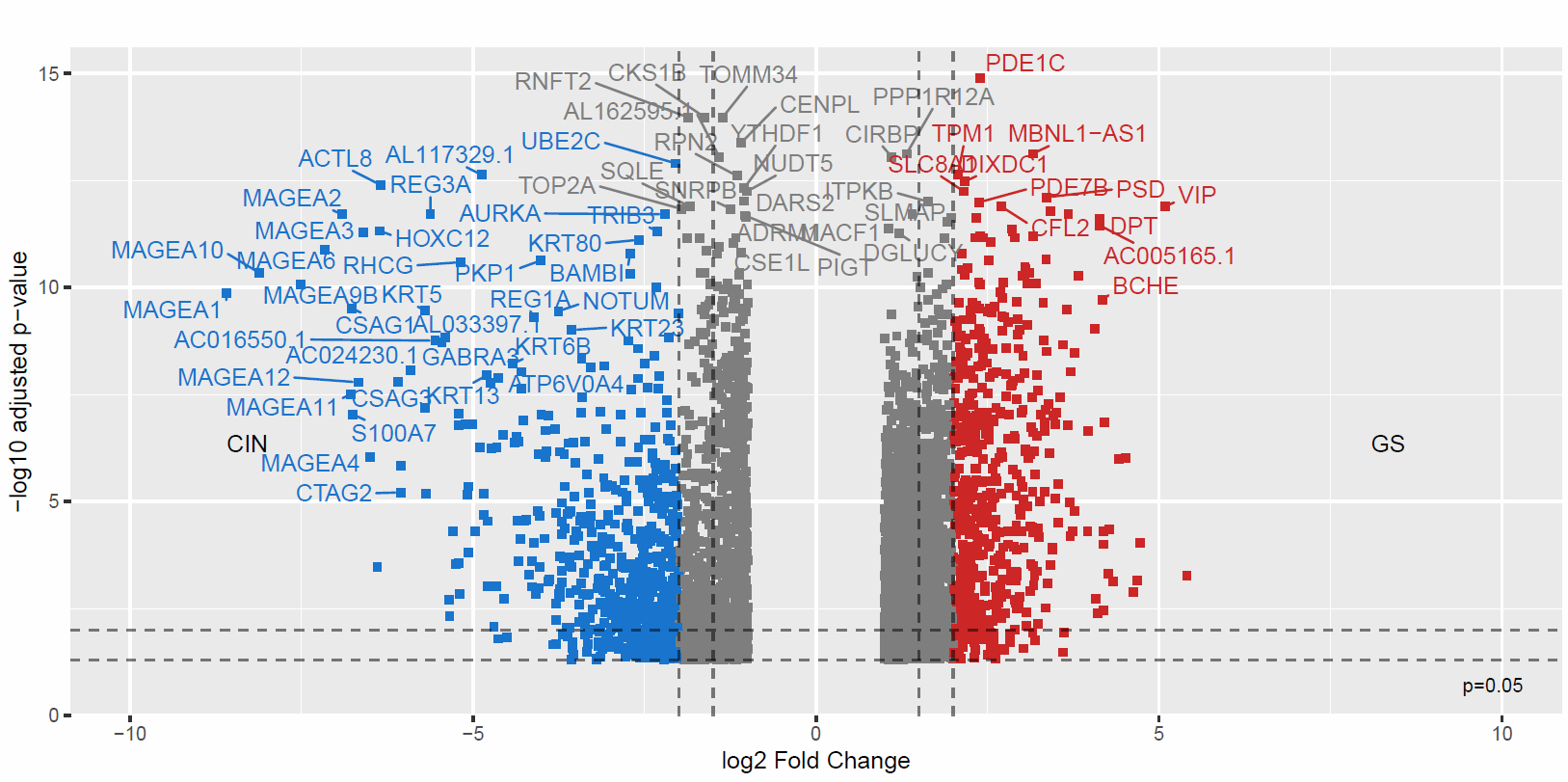 |

#### **Supplementary Figure 4c** Volcano plot of differential gene expressions between GTEx normal peritoneal (n-541) and liver (n=226) samples– 333 differentially expressed genes (adjusted p-value <0.05 & |log2FC| >2) were identified.

#### **Supplementary Figure 4d** Volcano plot of differential gene expressions between CIN (n=99) and GS (n=87) primary gastric cancer samples from the prospective cohort – 1172 differentially expressed genes (adjusted p-value <0.05 & |log2FC| >2) were identified. Samples were retrieved from the TCGA database.

Abbreviations: AMC, Academic Medical Center; PCA, principle component analysis; var., variance; FC, fold change

| **a** | **b** |
| --- | --- |
| 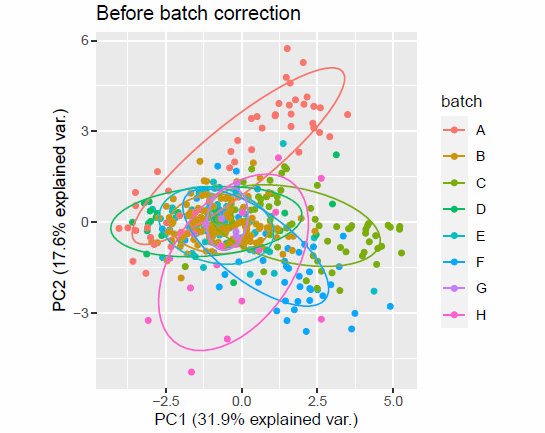 | 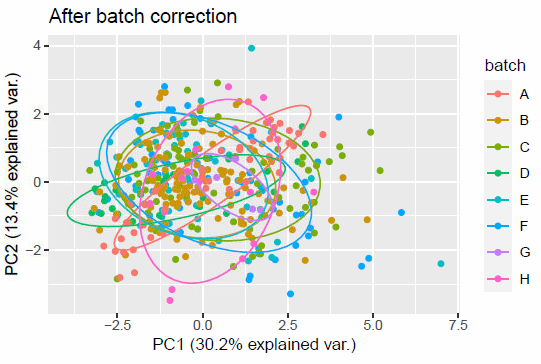 |

#### **Supplementary Figure 5a, b** Batch correction and PCA evaluation of gastric cancer DSP data

Abbreviations: DSP, digital spatial profiling; PCA, principle component analysis; var., variance

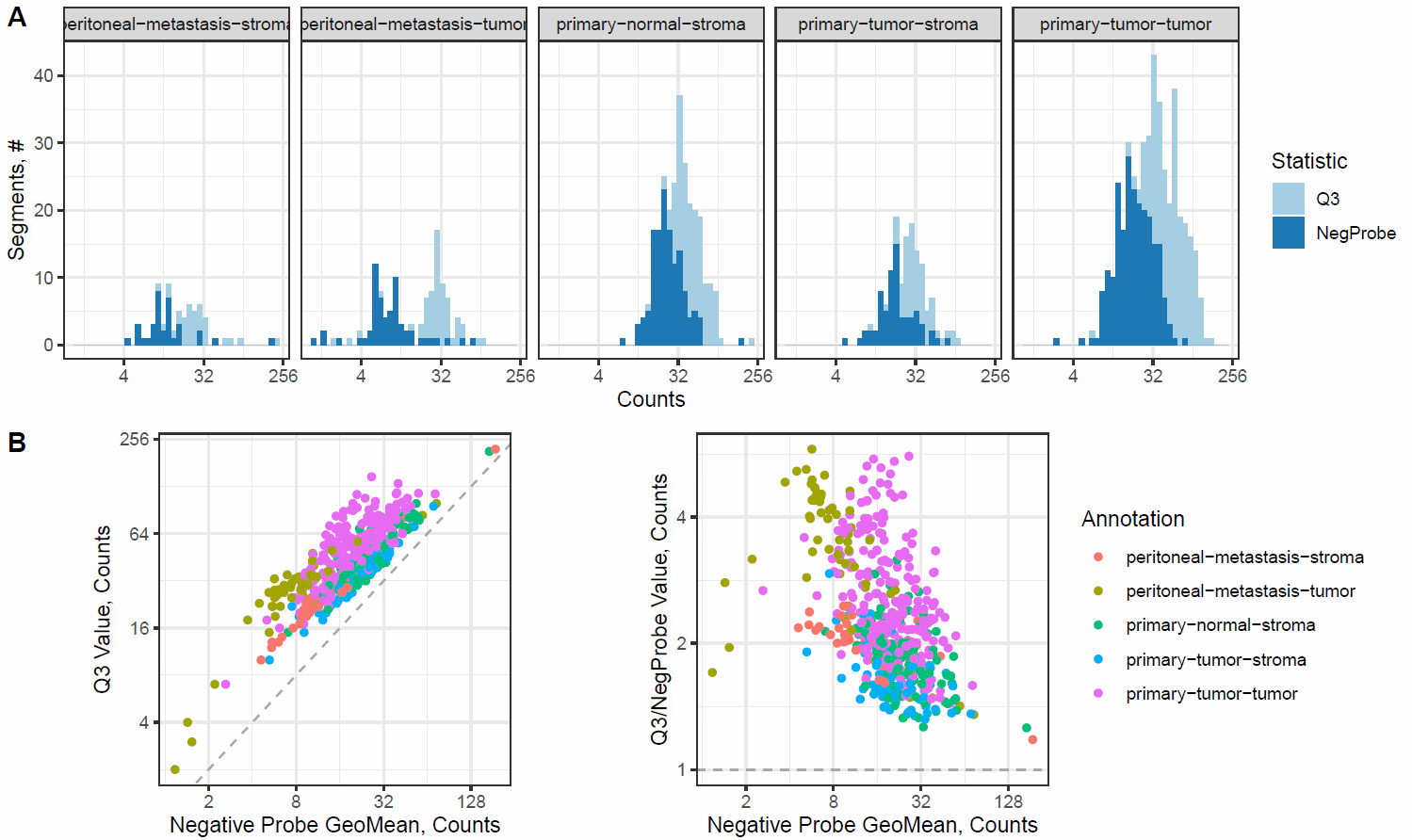

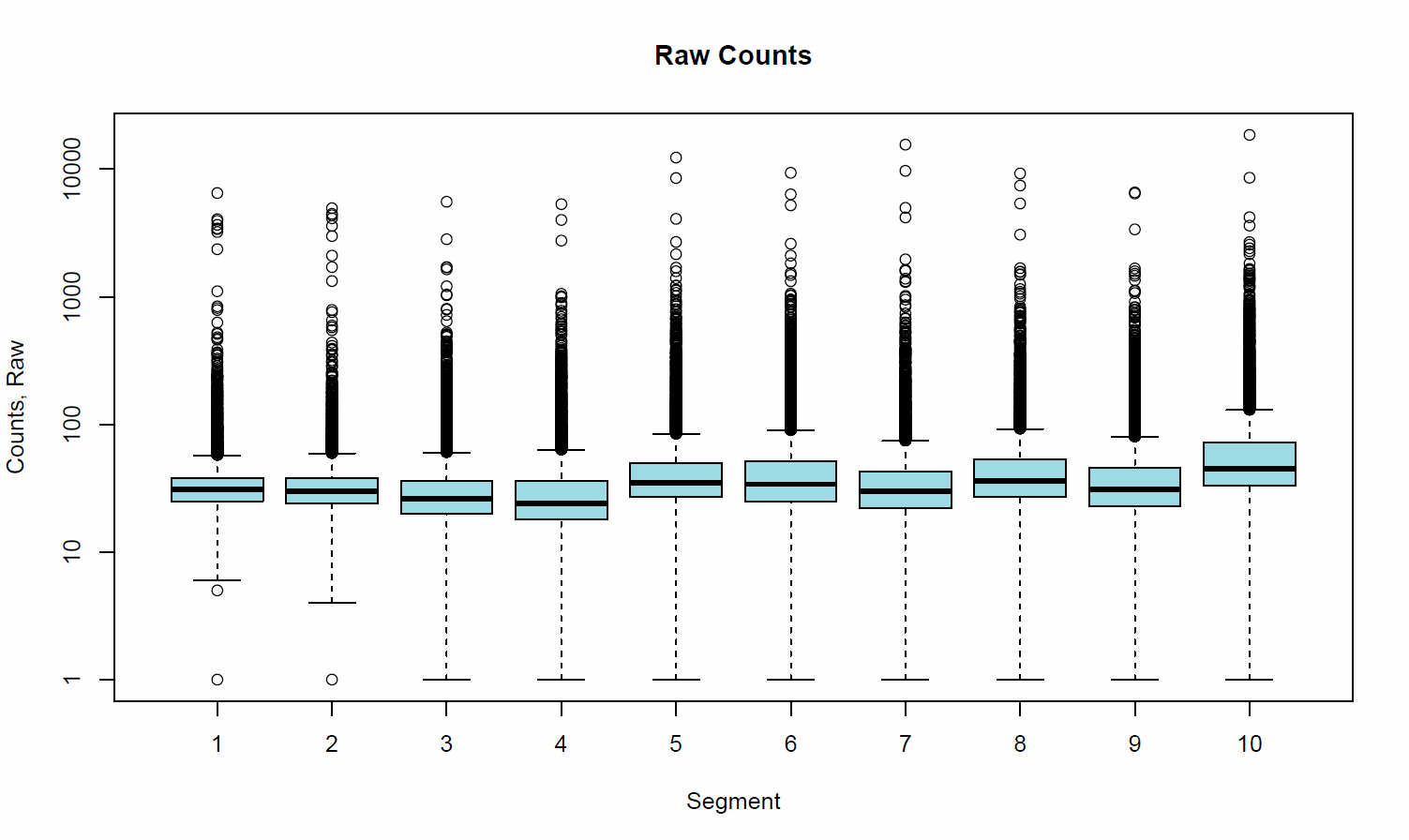

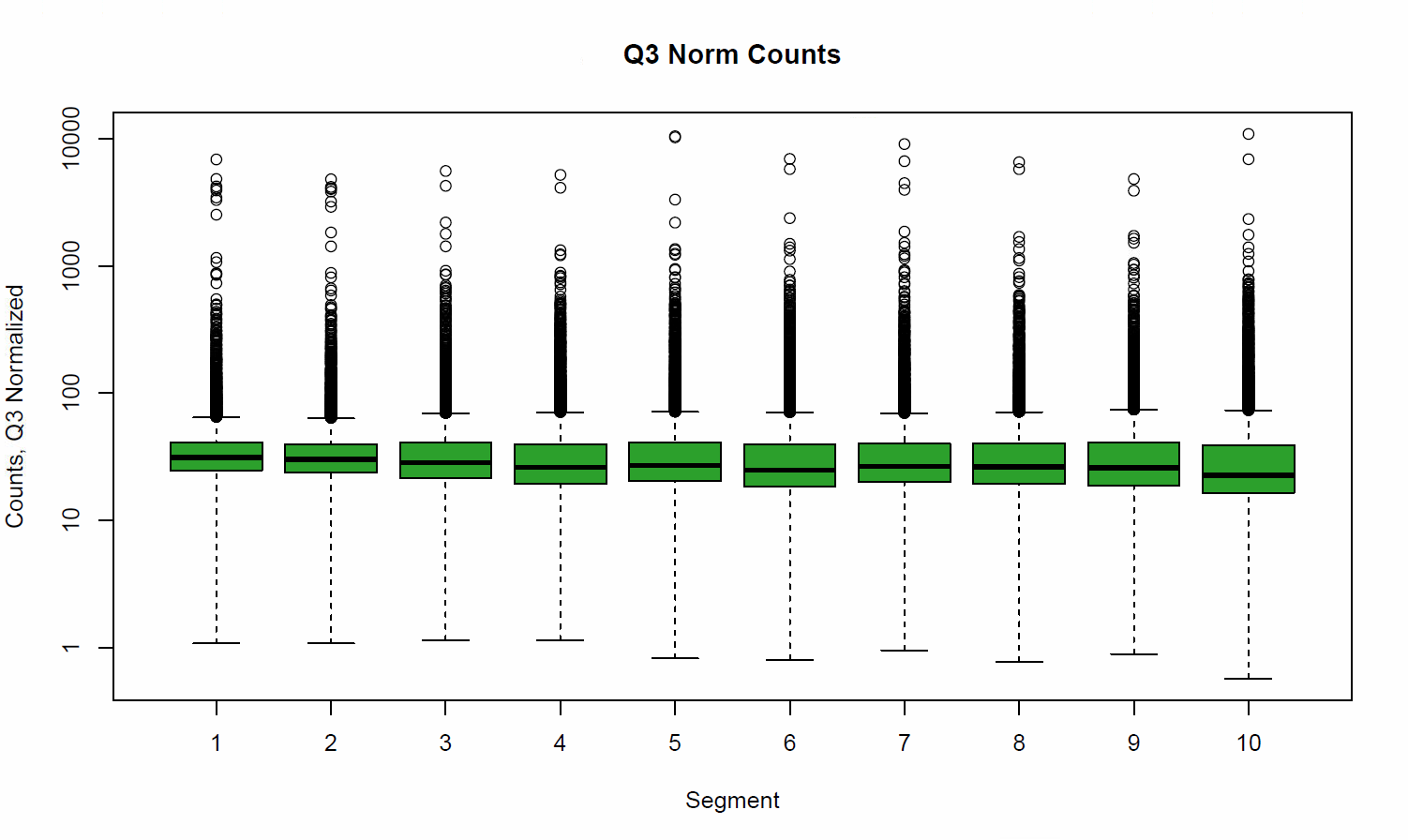

#### **Supplementary Figure 5c** Normalization evaluation of gastric cancer spatial transcriptomic data

| **d** | **e** |
| --- | --- |
| 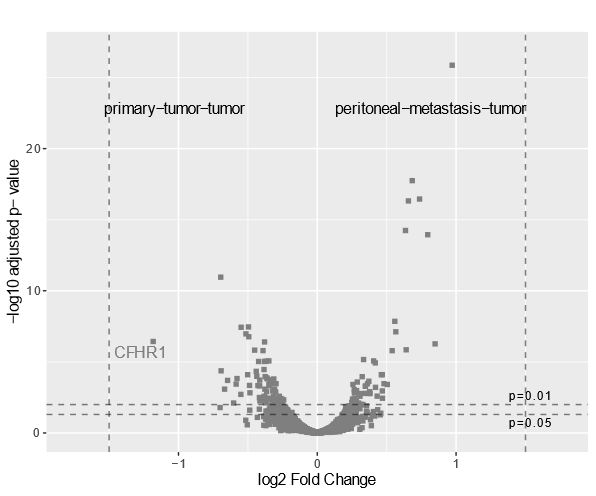 | 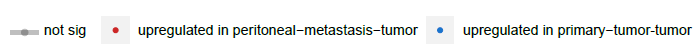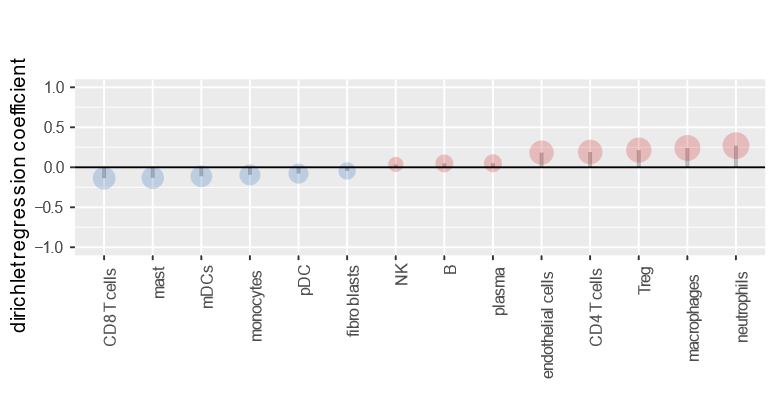 |
| **f** | **g** |
| 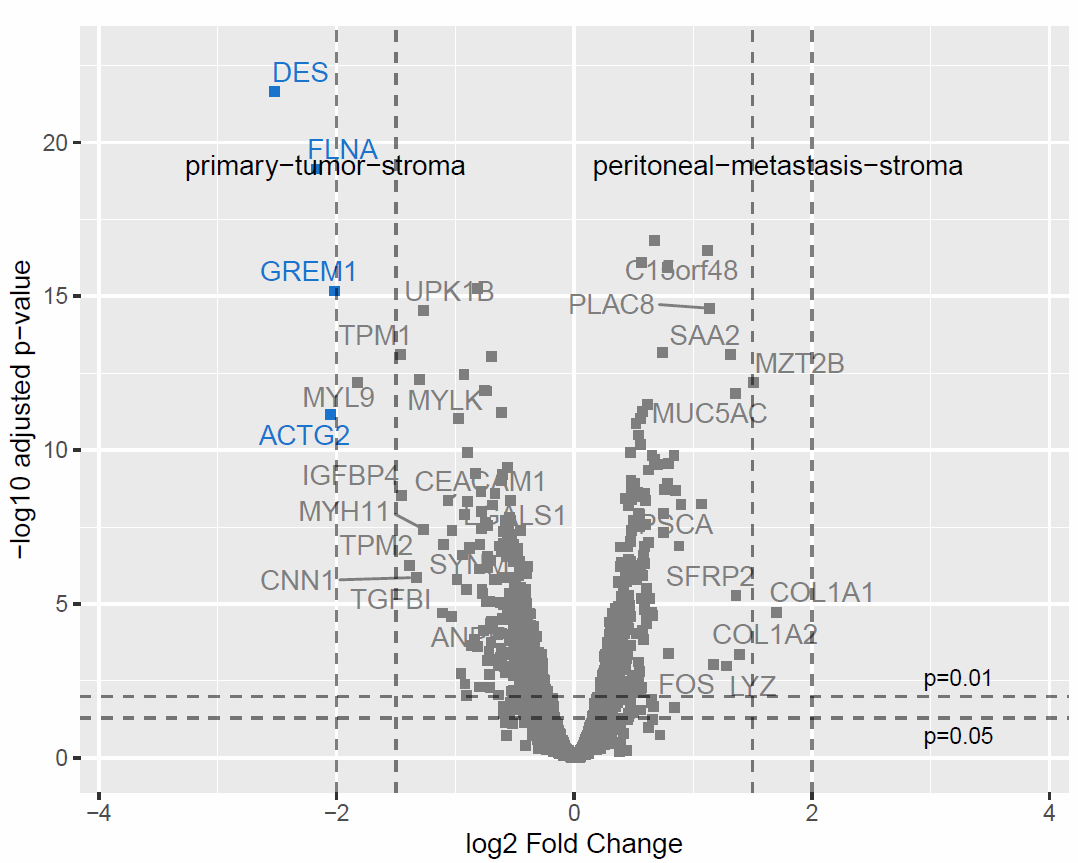 | 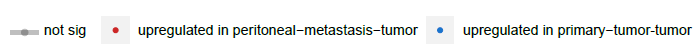  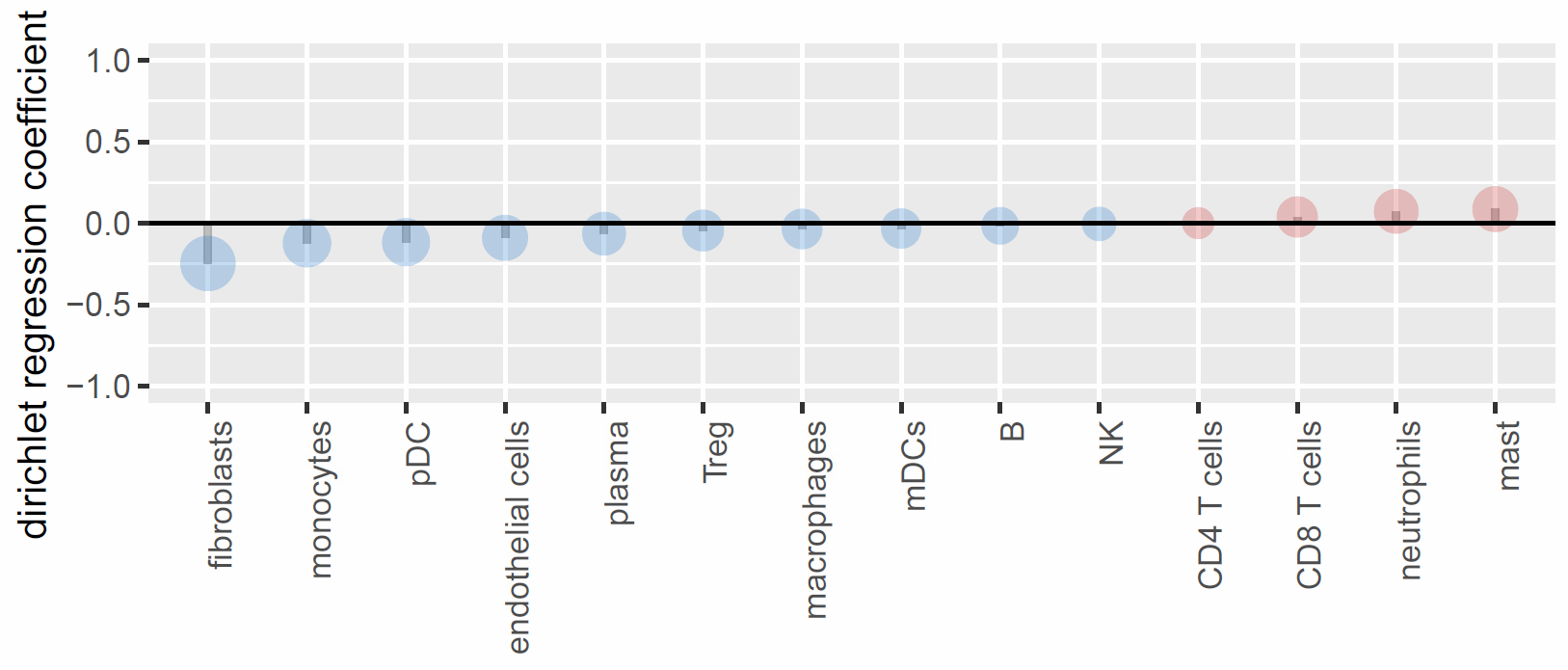 |

#### **Supplementary Figure 5d** Comparisons of differential gene expression in primary tumor tumor and peritoneal metastasis tumor

#### **Supplementary Figure 5e** Comparisons of immune cell proportion in primary tumor tumor and peritoneal metastasis tumor

#### **Supplementary Figure 5f** Comparisons of differential gene expression in primary tumor stroma and peritoneal metastasis stroma

#### **Supplementary Figure 5g** Comparisons of immune cell proportion in primary tumor stroma and peritoneal metastasis stroma

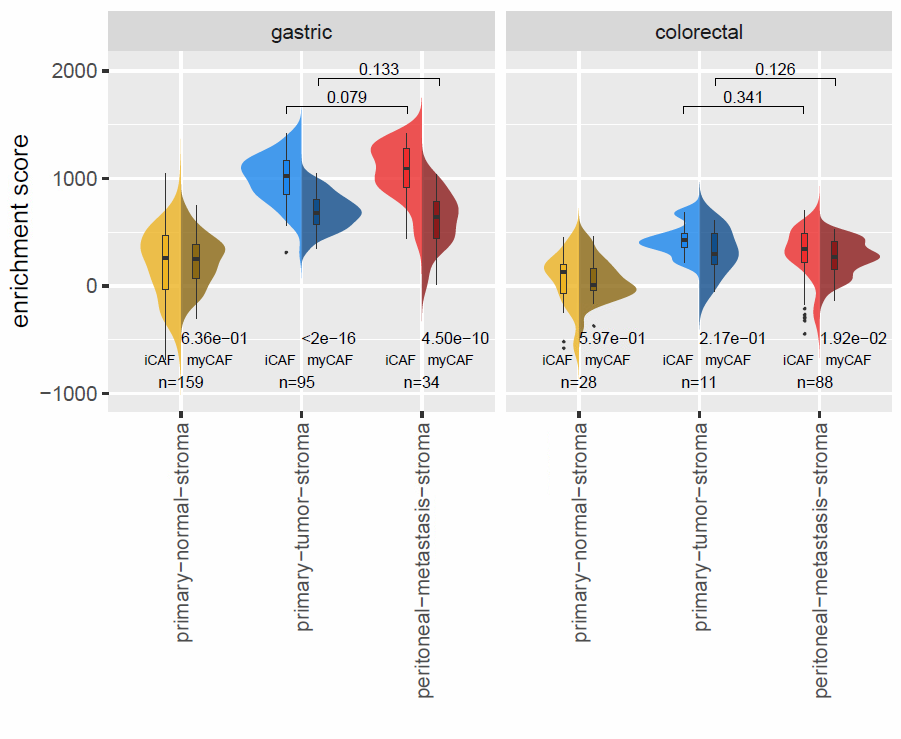

#### **Supplementary Figure 5h** Comparisons of ssGSEA enrichment scores of cancer associated fibroblasts subtypes in primary tumor stroma, peritoneal metastasis stroma and primary adjnormal stroma

Remarks: P-values were retrieved with a two-sided Wilcoxon test.

Abbreviations: n, number of regions of interest; iCAF, inflammatory cancer associated fibrobasts; myCAF, myofibroblast-like cancer associated fibroblasts; adjnormal, adjacent normal

**i**

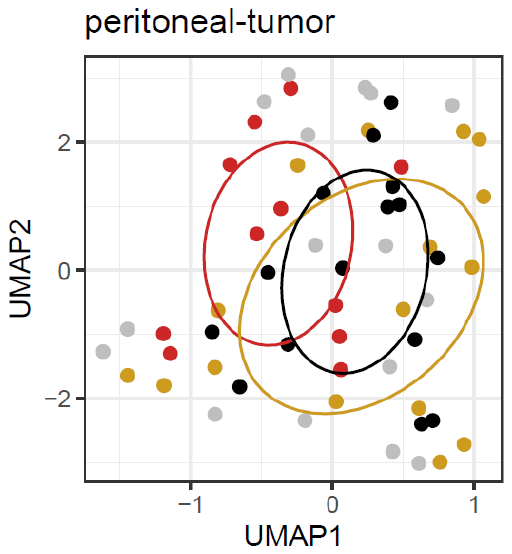

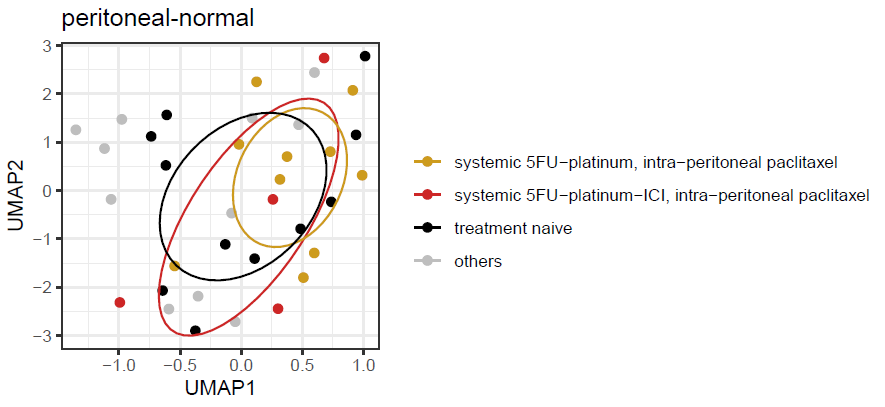

**j**

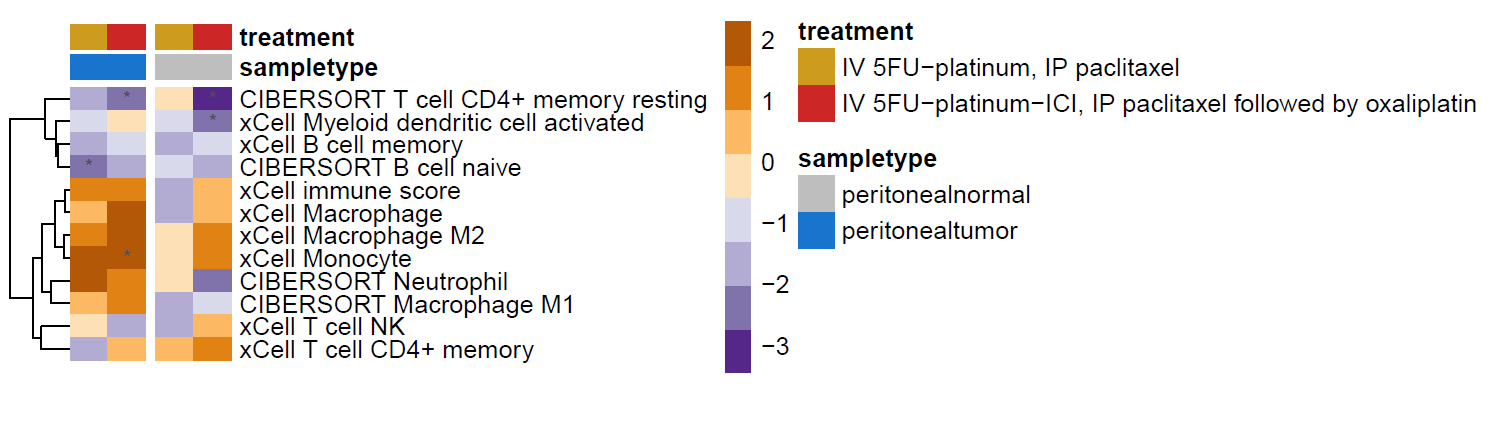

#### **Supplementary Figure 5i** UMAP of peritoneal-tumor and peritoneal-normal samples stratified by treatment status. **Supplementary Figure 5j** Heatmap of immune cell type differences against treatment naïve samples.

Heatmap values refer to GSEA NES for pathway comparisons and t-statistics from the unpaired t-test for immune cell type comparisons respectively.

Remarks: An asterisk refers to p-value<0.05.

Abbreviations: NES, normalized enrichment score; GSEA, Gene Set Enrichment Analysis; IV intra-venous; 5FU, 5-fluorouracil; ICI, immune checkpoint inhibition; IP, intra-peritoneal.

| **a** | **b** |
| --- | --- |
| 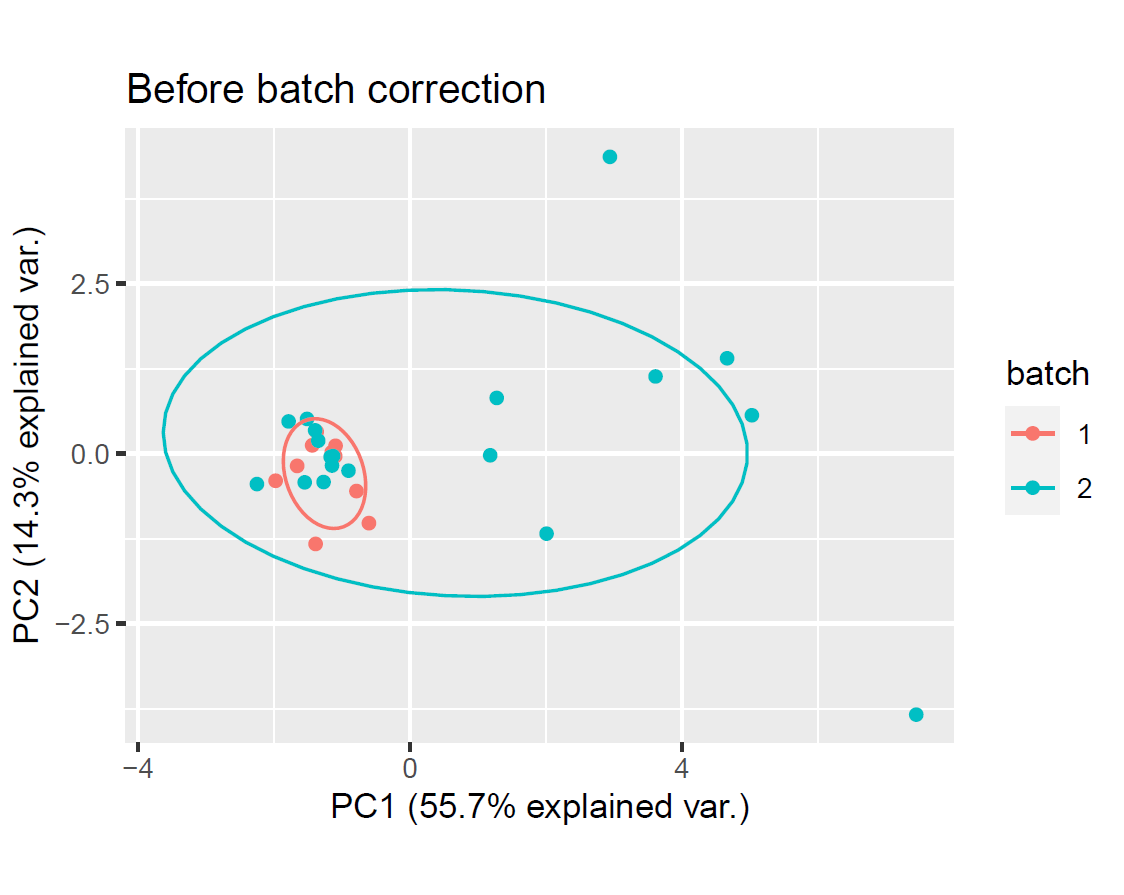 | 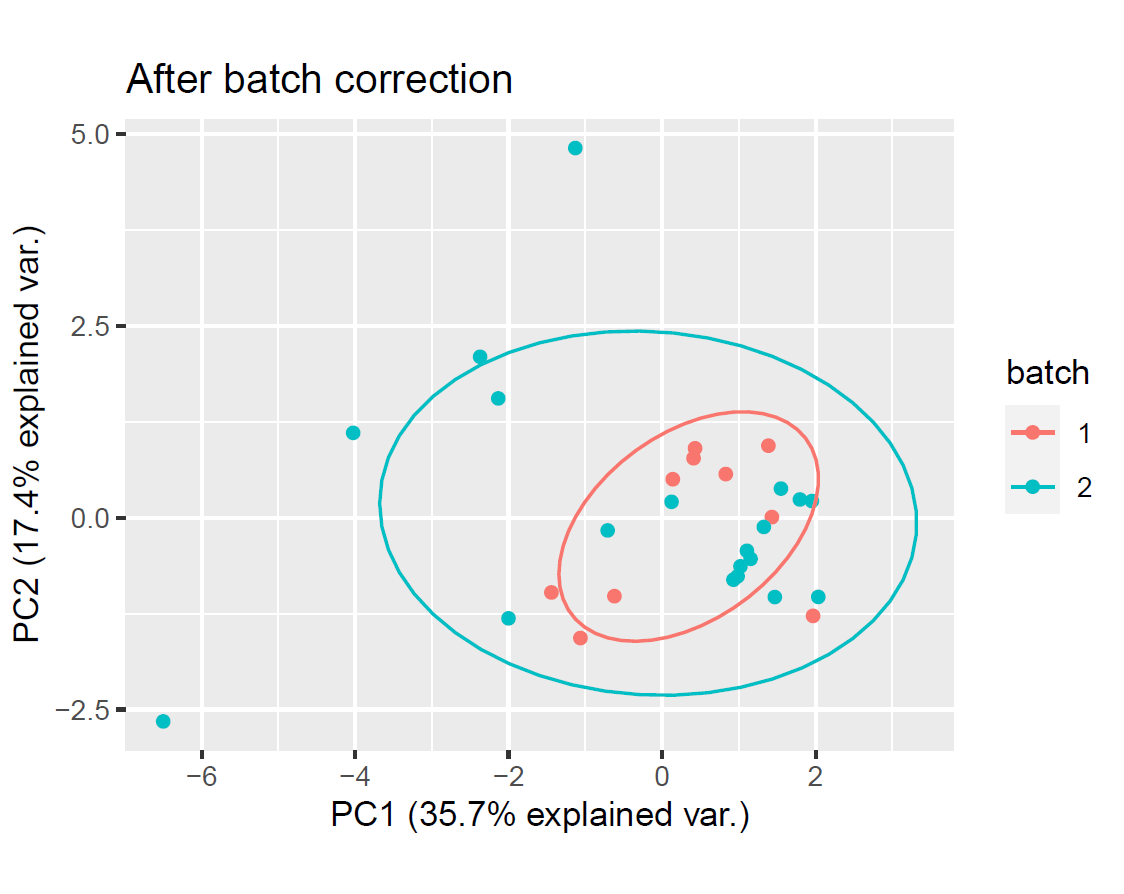 |

#### **Supplementary Figure 6a, b** Batch correction and PCA evaluation of Humice RNA-seq samples

Abbreviations: PC, principle component analysis; var., variance.

| **c** | **d** |
| --- | --- |
| 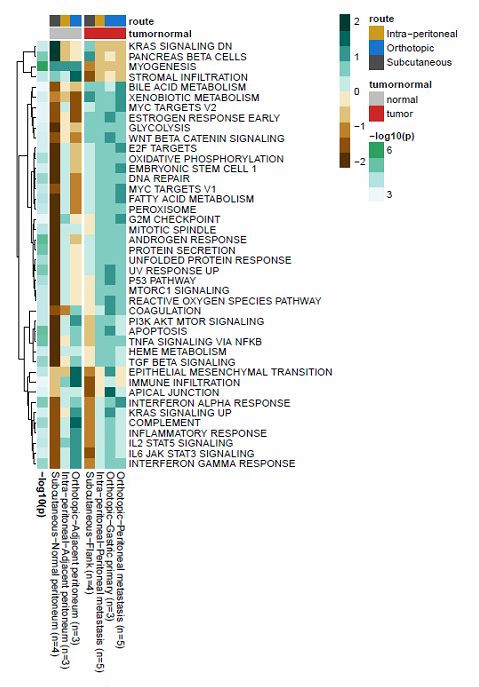 | 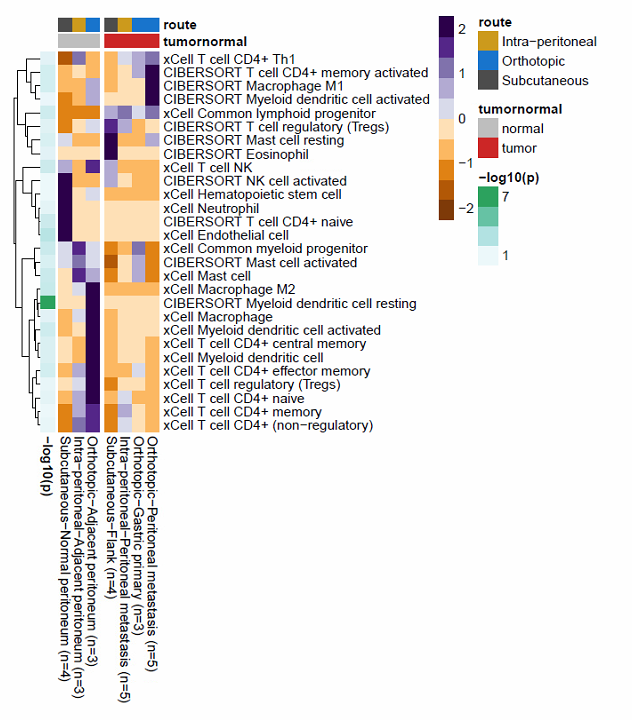 |

#### **Supplementary Figure 6c** Comparison of mean ssGSEA enrichment scores across sample types

#### **Supplementary Figure 6d** Comparison of mean cell type enrichment scores across sample types

Remarks: P values were retrieved with the Anova’s test.

Abbreviation: n=, number of samples; ssGSEA, single sample gene set enrichment analysis.
