## Supplementary Tables for "Spatially resolved niche and tumor microenvironmental alterations in gastric cancer peritoneal metastases"

**SUPPLMENTARY TABLES**

**CONTENT PAGE**

Remarks: Tables were numbers according to the relevant figures it was referenced to in the main manuscript. For example, the content of **Supplementary** **Table 1** is relevant to main **Figure 1**.

### SUPPLEMENTARY TABLES

|  |  | **Overall** | **no PM** | **metachronous PM** | **synchronous PM** | **p** |
| --- | --- | --- | --- | --- | --- | --- |
| **n** |  | **303** | **194** | **51** | **58** |  |
| **Age (median [IQR])** |  | 69.00 [60.00, 76.00] | 71.00 [63.22, 77.00] | 67.00 [58.50, 73.00] | 62.00 [56.00, 71.00] | <0.001 |
| **Gender (%)** | **Female** | 127 (42.8) | 77 ( 39.7) | 22 ( 43.1) | 28 ( 53.8) | 0.186 |
|  | **Male** | 170 (57.2) | 117 ( 60.3) | 29 ( 56.9) | 24 ( 46.2) |  |
| **Race (%)** | **Chinese** | 256 (85.9) | 173 ( 89.2) | 44 ( 86.3) | 39 ( 73.6) | 0.046 |
|  | **Indian** | 14 ( 4.7) | 5 ( 2.6) | 3 ( 5.9) | 6 ( 11.3) |  |
|  | **Malay** | 8 ( 2.7) | 5 ( 2.6) | 2 ( 3.9) | 1 ( 1.9) |  |
|  | **Others** | 20 ( 6.7) | 11 ( 5.7) | 2 ( 3.9) | 7 ( 13.2) |  |
| **Histology (%)** | **Adenocarcinoma** | 299 (98.7) | 194 (100.0) | 49 ( 96.1) | 56 ( 96.6) | 0.027 |
|  | **Others** | 4 ( 1.3) | 0 ( 0.0) | 2 ( 3.9) | 2 ( 3.4) |  |
| **Lauren classification (%)** | **Diffuse** | 77 (27.5) | 42 ( 21.9) | 20 ( 40.8) | 15 ( 38.5) | <0.001 |
|  | **Intestinal** | 132 (47.1) | 110 ( 57.3) | 19 ( 38.8) | 3 ( 7.7) |  |
|  | **Mixed** | 37 (13.2) | 27 ( 14.1) | 6 ( 12.2) | 4 ( 10.3) |  |
|  | **NOS** | 34 (12.1) | 13 ( 6.8) | 4 ( 8.2) | 17 ( 43.6) |  |
| **T stage (%)** | **T1** | 32 (12.0) | 32 ( 16.5) | 0 ( 0.0) | 0 ( 0.0) | <0.001 |
|  | **T2** | 22 ( 8.3) | 22 ( 11.3) | 0 ( 0.0) | 0 ( 0.0) |  |
|  | **T3** | 90 (33.8) | 69 ( 35.6) | 13 ( 26.5) | 8 ( 34.8) |  |
|  | **T4** | 122 (45.9) | 71 ( 36.6) | 36 ( 73.5) | 15 ( 65.2) |  |
| **N stage (%)** | **N0** | 63 (24.0) | 59 ( 30.4) | 1 ( 2.1) | 3 ( 14.3) | <0.001 |
|  | **N1** | 40 (15.2) | 34 ( 17.5) | 5 ( 10.4) | 1 ( 4.8) |  |
|  | **N2** | 49 (18.6) | 35 ( 18.0) | 8 ( 16.7) | 6 ( 28.6) |  |
|  | **N3** | 111 (42.2) | 66 ( 34.0) | 34 ( 70.8) | 11 ( 52.4) |  |

#### **Supplementary Table 1a** Overview of cohort

Remarks: P-values were derived from a two-sided Fisher’s exact test for categorical variables and two-sided Wilcoxon test for continuous variables.

Abbreviations: PM, peritoneal metastasis; IQR, interquartile range; CIN, chromosomal instability; GS genomically stable; MSI, microsatellite instability; EBV, Epstein-Barr virus; NOS, not otherwise specified

|  | **primary-tumor** | **peritoneal-adjnormal** | **peritoneal-tumor** | **primary- adjnormal** |
| --- | --- | --- | --- | --- |
| **primary-tumor** | - | N=14, n=36 | N=16, n=46 | N=10, n=21 |
| **peritoneal-normal** | N=14, n=36 | - | N=35, n=123 | N=11, n=28 |
| **peritoneal-tumor** | N=16, n=46 | N=35, n=123 | - | N=12, n=32 |
| **primary-normal** | N=10, n=21 | N=11, n=28 | N=12, n=32 | - |

#### **Supplementary Table 1b** Overview of paired bulk RNA-seq gastric cancer samples

Abbreviations: N=, number of patients; n=, number of samples; adjnormal, adjacent normal

|  |  | **Overall** | **primary-tumor-no PM** | **primary-tumor-with PM** | **peritoneal-tumor** | **liver-tumor** | **p** |
| --- | --- | --- | --- | --- | --- | --- | --- |
| **Number of samples** |  | **256** | **94** | **49** | **100** | **13** |  |
| **EMT score (Tagliazucchi *et al.*) (mean (SD))** |  | 0.02 (1.25) | -0.25 (1.45) | -0.28 (1.23) | 0.45 (0.97) | -0.24 (0.72) | <0.001 |
| **Tumor purity (ESTIMATE) (mean (SD))** |  | 0.60 (0.16) | 0.66 (0.17) | 0.59 (0.16) | 0.55 (0.13) | 0.66 (0.17) | <0.001 |
| **TME subtype (Bagaev *et al.*) (%)** | **Deserted** | 87 (34.0) | 45 ( 47.9) | 15 ( 30.6) | 21 ( 21.0) | 6 ( 46.2) | 0.004 |
|  | **Fibrotic** | 63 (24.6) | 14 ( 14.9) | 12 ( 24.5) | 35 ( 35.0) | 2 ( 15.4) |  |
|  | **Immune-enriched** | 59 (23.0) | 23 ( 24.5) | 12 ( 24.5) | 21 ( 21.0) | 3 ( 23.1) |  |
|  | **Immune-enriched, fibrotic** | 47 (18.4) | 12 ( 12.8) | 10 ( 20.4) | 23 ( 23.0) | 2 ( 15.4) |  |
| **Mesenchymal GC (Ho *et al.*) (%)** | **MesGC** | 115 (47.7) | 22 ( 25.0) | 21 ( 43.8) | 67 ( 72.8) | 5 ( 38.5) | <0.001 |
|  | **nonMesGC** | 126 (52.3) | 66 ( 75.0) | 27 ( 56.2) | 25 ( 27.2) | 8 ( 61.5) |  |
| **GI tumor lineage (Wang *et al.*) (%)** | **Gastric-dominant** | 132 (51.6) | 57 ( 60.6) | 19 ( 38.8) | 45 ( 45.0) | 11 ( 84.6) | 0.003 |
|  | **GI-mixed** | 124 (48.4) | 37 ( 39.4) | 30 ( 61.2) | 55 ( 55.0) | 2 ( 15.4) |  |
| **Lauren classification (%)** | **Diffuse** | 60 (30.2) | 17 ( 18.1) | 14 ( 33.3) | 29 ( 46.0) |  | <0.001 |
|  | **Intestinal** | 76 (38.2) | 56 ( 59.6) | 10 ( 23.8) | 10 ( 15.9) |  |  |
|  | **Mixed** | 18 ( 9.0) | 11 ( 11.7) | 7 ( 16.7) | 0 ( 0.0) |  |  |
|  | **NOS** | 45 (22.6) | 10 ( 10.6) | 11 ( 26.2) | 24 ( 38.1) |  |  |
| **TCGA subtype (%)** | **CIN** | 81 (48.2) | 62 ( 66.0) | 15 ( 41.7) | 4 ( 10.5) |  | <0.001 |
|  | **EBV** | 6 ( 3.6) | 4 ( 4.3) | 1 ( 2.8) | 1 ( 2.6) |  |  |
|  | **GS** | 69 (41.1) | 17 ( 18.1) | 19 ( 52.8) | 33 ( 86.8) |  |  |
|  | **MSI** | 12 ( 7.1) | 11 ( 11.7) | 1 ( 2.8) | 0 ( 0.0) |  |  |
| **Fraction of altered genome (CNV) (mean (SD))** |  | 0.21 (0.23) | 0.29 (0.24) | 0.17 (0.22) | 0.05 (0.11) |  | <0.001 |
| **Whole genome duplication (%)** | **No** | 88 (60.3) | 48 ( 51.1) | 24 ( 70.6) | 16 ( 88.9) |  | 0.0045 |
|  | **Yes** | 58 (39.7) | 46 ( 48.9) | 10 ( 29.4) | 2 ( 11.1) |  |  |
| **Clonality (%)** | **Monoclonal** | 59 (42.4) | 33 ( 36.7) | 16 ( 48.5) | 10 ( 62.5) |  | 0.119 |
|  | **Polyclonal** | 80 (57.6) | 57 ( 63.3) | 17 ( 51.5) | 6 ( 37.5) |  |  |
| **Number of clones (median [IQR])** |  | 2.00 [1.00, 3.00] | 2.00 [1.00, 3.00] | 2.00 [1.00, 2.00] | 1.00 [1.00, 2.00] |  | 0.005 |

#### **Supplementary Table 4a** PT-PM differences in GCPM

Remarks: EMT scores were derived by methods described by Tagliazucchi et al^1^. Derivation of gastric-dominant vs GI-mixed subtype was conducted using a 12-gene signature reported by Wang et al^2^ in the original article, with steps reported in the article’s Supplementary Figure 24. For each sample, a score of 1 or -1 for each of the 12-signature gene was allocated based on its relative expression (> or ≤ median) and whether the gene is associated with the Gastric-dominant or GI-mixed feature. Signature scores per patient were summed up and subsequently classified into either a GI-mixed (signature score >median) or Gastric-dominant (signature score ≤median) subgroup. MesGC subtypes were retrieved using the NTP algorithm as reported by Ho et al^3^. TME subtypes were retrieved utilizing methods described by Bagaev et al^4^. Tumor purity was derived utilizing the ESTIMATE algorithm as described by Yoshihara et al^5^.

Abbreviations: MesGC, gastric cancer mesenchymal subtype; nonMesGC, gastric cancer non-mesenchymal subtype; GC, gastric cancer; NTP, nearest template prediction; CIN, chromosomal instability; GS genomically stable; MSI, microsatellite instability; EBV, Epstein-Barr virus; TME, tumor microenvironment; IQR, inter-quartile range.

|  | **Pass** | **Warning** |
| --- | --- | --- |
| **Low Reads** | 712 | 1 |
| **Low Trimmed** | 713 | 0 |
| **Low Stitched** | 683 | 30 |
| **Low Aligned** | 669 | 44 |
| **Low Saturation** | 707 | 6 |
| **Low Negatives** | 713 | 0 |
| **Low Nuclei** | 713 | 0 |
| **Low Area** | 663 | 50 |
| **TOTAL FLAGS** | **712** | 1 |

#### **Supplementary Table 5a** Quality evaluation and inclusion of DSP ROIs

Abbreviations: DSP, Digital spatial profiling; ROI, region of interest.

| **ROI** | **primary-adjnormal (n)** | **primary-tumor (n)** | **peritoneal-metastasis (n)** |
| --- | --- | --- | --- |
| **stroma** | 159 | 95 | 34 |
| **tumor** | 0 | 314 | 60 |

#### **Supplementary Table 5b** Distribution of ROIs in spatial transcriptomic analysis of gastric cancer samples

Abbreviations: n, number of regions of interest; ROI, regions of interest; adjnormal, adjacent normal

| **Reference** | **Name of gene set/signature** | **Remarks** |
| --- | --- | --- |
| **Liberzon *et al.*^6^** | Molecular Signatures Database (MSigDB) hallmark gene set collection |  |
| **Jiang *et al.*^7^** | T cell dysfunction: T cell exhaustion; M2/M1 TAM |  |
| **Öhlund *et al.*^8^** | Inflammatory CAF, myofibroblastic-like CAF | Genes with log 2 fold change value greater than 2.5, adjusted p-value <0.05 were included |
| **Yoshihara *et al*.^5^** | Stromal score, Immune score – Estimate algorithm |  |
| **Ben-Porath *et al.*^9^** | Embryonic stem cell signature 1 & 2 |  |
| **Wang *et al*.^10^** | YAP-TAZ/Hippo pathway |  |
| **Kieffer *et al.^11^*** | CAF subtype signatures |  |
| **Wang *et al*.^2^** | GI-mixed vs gastric-dominant subtype | Implementation and assignment of GI-mixed vs gastric-dominant subtype was in concordance with methods described in the Supplementary Figure 24 of Wang et al. |
| **Ho *et al.*^12^** | Mesenchymal subtype for gastric cancer (Mes-GC) | Implementation and assignment of the Mes-GC subtype was conducted with the nearest template prediction algorithm |
|  | Selected genes utilized for target gene analysis | Tumor related genes: *APC, BRAF, CLDN18 (and CLDN18.2 isoform), CTNNB1, DKK1, EGFR, ERBB2, CAPRIN1, FBXW7, FGFR1, FGFR2, KEAP1, MET, MLH1, MYC, NFE2L2, NOTCH1, NTRK1, NTRK2, NTRK3, PIK3CA, PIK3R1, RHOA,*  *SMAD4, STK11, TACSTD2, TEAD1, TEAD2, TEAD3, TEAD4*  Immune related genes: *CD274, CD8A, CTLA4, FOXP3, GZMB, HAVCR2, IDO1,*  *IDO2, IFNG, LAG3, PDCD1, PRF1, TIGIT* |

#### **Supplementary Table** **6** Gene signature sets

Abbreviations: CAF, cancer associated fibroblasts; GI, gastro-intestinal
